# Broadly Neutralizing Antibody Epitopes on HIV-1 Particles are exposed after Virus Interaction with Host Cells

**DOI:** 10.1101/2023.01.20.524996

**Authors:** Priyanka Gadam Rao, Gregory S. Lambert, Chitra Upadhyay

**Affiliations:** Division of Infectious Disease, Department of Medicine, Icahn School of Medicine at Mount Sinai, New York, New York, USA

**Keywords:** Human immunodeficiency virus, neutralization, antibodies, broadly neutralizing antibodies (bNAbs), non-neutralizing antibodies (nNAbs), envelope conformations, glycosylation, lectins, glycans.

## Abstract

The envelope glycoproteins (Env) on HIV-1 virions are the sole target of broadly neutralizing antibodies (bNAb) and the focus of vaccines. However, many cross-reactive conserved epitopes are often occluded on virus particles, contributing to the evasion of humoral immunity. This study aimed to identify the Env epitopes that are exposed/occluded on HIV-1 particles and to investigate the mechanisms contributing to their masking. Using a flow cytometry-based assay, three HIV-1 isolates, and a panel of antibodies, we show that only select epitopes including V2i, gp120-g41 interface, and gp41-MPER are accessible on HIV-1 particles, while V3, V2q, and select CD4bs epitopes are masked. These epitopes become accessible after allosteric conformational changes are induced by pre-binding of select Abs, prompting us to test if similar conformational changes are required for these Abs to exhibit their neutralization capability. We tested HIV-1 neutralization where virus-mAb mix was pre-incubated/not pre-incubated for one hour prior to adding the target cells. Similar levels of neutralization were observed under both assay conditions, suggesting that the interaction between virus and target cells sensitizes the virions for neutralization via bNAbs. We further show that lectin-glycan interactions can also expose these epitopes. However, this effect is dependent on the lectin specificity. Given that, bNAbs are the ideal for providing sterilizing immunity and are the goal of current HIV-1 vaccine efforts, these data offer insight on how HIV-1 may occlude these vulnerable epitopes from the host immune response. In addition, the findings can guide the formulation of effective antibody combinations for therapeutic use.

**Importance:** The human immunodeficiency virus (HIV-1) envelope (Env) glycoprotein mediates viral entry, and is the sole target of neutralizing antibodies. Our data suggest that antibody epitopes including V2q (e.g., PG9, PGT145), CD4bs (e.g., VRC01, 3BNC117) and V3 (2219, 2557) are masked on HIV-1 particles. The PG9 and 2219 epitopes became accessible for binding after conformational unmasking was induced by pre-binding of select mAbs. Attempts to understand the masking mechanism led to the revelation that interaction between virus and host cells is needed to sensitize the virions for neutralization by broadly neutralizing antibodies (bNAbs). These data provide insight on how bNAbs may gain access to these occluded epitopes to exert their neutralization effects and block HIV-1 infection. These findings have important implications for the way we evaluate the neutralizing efficacy of antibodies and can potentially guide vaccine design.

## 1. Introduction

The envelope glycoprotein (Env) of human immunodeficiency virus type-1 (HIV-1) is the virus attachment protein that interacts with the host-cell receptor CD4 and chemokine receptors CCR5 or CXCR4 to initiate infection (1). HIV-1 displays an average of 7-14 Env trimers per particle. However, this number varies among isolates (2). Env is synthesized as a gp160 precursor protein in the endoplasmic reticulum (ER), where signal peptide cleavage, folding, addition of high-mannose glycans, and trimerization in association with molecular chaperones takes place (3–6). Once the nascent polypeptide attains its native folding state, the Env egresses from the ER and translocates to the Golgi apparatus (3, 7–13). In the Golgi, gp160 is cleaved by host furin-like proteases to generate a transmembrane gp41 subunit and a non-covalently associated surface gp120 subunit. Three gp120-gp41 heterodimers assemble to form the trimeric functional Env spikes that are then directed to the plasma membrane for incorporation into virions (3, 11–13). The gp120 subunit comprises five variable (V1-V5) and five conserved (C1-C5) regions. The V1V2 from each of the protomers join at the top to form the trimer apex, an immunogenic, structurally conserved region targeted by some of the most potent HIV-1 bNAbs (14–18). The gp41 subunit comprises the fusion machinery, mediating entry of the virus into the target cell by facilitating membrane fusion (19, 20). While trimeric Env is expected to be the most abundant form of Env present on virions, non-functional Env such as dimers, monomers, and gp41 stumps may also be present (21–23). These nonfunctional Env are immunodominant and act as decoys to divert the immune response away from vulnerable conserved epitopes resulting in higher titers of non-neutralizing antibody responses (23, 24), further adding a layer of diversity to the already complex Env protein.

HIV-1 Env is the only viral protein on the surface of HIV-1 particles, and is therefore a principal target for HIV-1 broadly neutralizing antibodies (bNAbs) that neutralize the virus and prevent infection of host cells. Passive transfer of various bNAbs has been shown to protect non-human primates and humanized mice from viral challenge (25, 26). Thus, bNAbs are being investigated for both therapeutic use and to guide the design of Env-based immunogens (27, 28). Major efforts are being invested in improving HIV-1 antigen design and strategies that can stimulate production of antibodies with similarly broad and neutralizing responses. Several approaches using the trimer-mimic and virus-like particles have been explored, but eliciting bNAbs via vaccination has proven extremely difficult. Despite large research efforts, immunogens capable of eliciting bNAbs do not exist yet. HIV-1 has evolved multiple mechanisms such as high antigenic diversity, conformational flexibility, and a dense coat of N-linked glycans that masks the conserved epitopes on Env (29). This masking of conserved epitopes is one of several factors that has impeded the development of an Env-derived immunogen capable of eliciting bNAbs against HIV-1. Currently, five regions have been identified on the HIV-1 Env that are the focus of bNAb elicitation. These include the V2 apex, the V3-N332 supersite, the membrane proximal external region (MPER), the CD4 binding site (CD4bs), and the gp120-gp41 interface (30–32). Thus, identifying the Env epitopes that are occluded on virions and understanding the mechanisms that govern their unmasking is crucial to inform design and development of a protective HIV-1 vaccine.

Several monoclonal antibodies (mAbs) have been isolated from HIV-1 infected individuals and vaccinees from clinical trials that have helped to identify and map the vulnerable Env epitopes. These mAbs are categorized into distinct types based on their neutralization breadth and potency. The highly potent neutralizing antibodies that can neutralize broad arrays of HIV-1 isolates are referred as bNAbs. The weak- or non-neutralizing antibodies (wNAbs, nNAbs) bind to the Env in a manner that is not efficient at blocking virus infection. The bNAbs known to date have been isolated from HIV-1 infected individuals, and elicitation of these kind of mAbs via vaccination has not yet been achieved. In contrast, nNAbs or wNAbs, including those that target the V1V2 and V3 regions, are readily elicited via various Env vaccines as well as during HIV-1 infection (33–42). In fact, the early antibody responses are typically directed against the gp41 domain of Env and later evolves to include gp120 (43). Only after several years of infection do a fraction of infected individual develop bNAbs (44). The HIV-1 antibodies are further categorized based on the distinct Env region that they target, such as V1V2, V3, CD4bs, gp41-MPER, and the gp120-gp41 interface (45). While most mAbs are conformation-dependent, some also recognize linear epitopes. For example, the V1V2 mAbs are categorized into three types (V2i, V2p and V2q) based on their binding mode (46, 47). V2i-specific mAbs, (e.g., 697, 830A, and 2158) recognize V1V2 when its V2C region is in a β-strand configuration, as determined by X ray crystallography; the epitope region of these V2i mAbs is discontinuous, highly conformational, and overlaps the α4β7 integrin-binding motif (46). The V2p mAbs (eg., CH58 and CH59) were isolated from RV144 vaccinees and target a linear peptide region on V2; these mAbs recognize V2 when the V2C strand region is in an α-helix and extended coil conformation (48–50). The epitope region recognized by V2q mAbs (e.g., PG9, PG16, and PGT145) includes two N-linked glycans (N156 and N160) and consists of the V2C in its β-strand configuration (15, 51–53); only V2q mAbs are bNAbs. Another key distinction is whether the mAbs recognize Env in the context of functional trimers for example, PG9 and PG16 (21). The conformational diversity of Env, and the presence of functional and non-functional Envs on virions, can significantly influence both antibody efficacy and their elicitation.

In this study, we used a panel of Abs specific to different regions of Env to identify the overall antigenic makeup of Env on virus particles. We established a flow cytometry-based assay to evaluate the neutralizing and non-neutralizing epitopes that are masked or accessible on virions, and to identify the mechanisms that can unmask the occluded epitopes. We show that virus interaction with host cells induces changes in Env, exposing the epitopes and allowing antibody binding and virus neutralization. We further show that a virus-mAb pre-incubation step, widely used in neutralization assays in the field, does not appear to be required for neutralization by these bNAbs. Similar exposure of epitopes can also be achieved by lectin and Env glycan interactions. However, this activity is dependent on the lectin specificity. These findings have important implications for the way we evaluate the neutralizing efficacy of antibodies, and provides important clues for improving vaccine design and the formulation of effective antibody combinations for therapeutic use.

## 2. Materials and Methods

### Cell lines

HEK293T/17 cells were obtained from the American Type Culture Collection (ATCC, Manassas, VA). The following reagent was obtained through the NIH HIV Reagent Program, Division of AIDS, NIAID, NIH: TZM-bl Cells, ARP-8129, contributed by Dr. John C. Kappes, Dr. Xiaoyun Wu, and Tranzyme Inc. (54). For all experiments, HEK293T/17 cells (293T) were used to produce infectious HIV-1 viruses and the TZM.bl cell line was used to assay virus infectivity. TZM.bl cell line is derived from HeLa cells and is genetically modified to express high levels of CD4, CCR5 and CXCR4 and contains reporter cassettes of luciferase and β-galactosidase that are each expressed from an HIV-1 LTR. The 293T and TZM.bl cell lines were routinely sub-cultured every 3 to 4 days by trypsinization and were maintained in Dulbecco’s Modified Eagle’s Medium (DMEM) supplemented with 10% heat-inactivated fetal bovine serum (FBS), HEPES pH 7.4 (10 mM), L-glutamine (2 mM), penicillin (100 U/ml), and streptomycin (100 μg/ml) at 37°C in a humidified atmosphere with 5% CO2.

### Plasmids

A full-length transmitted/founder (T/F) infectious molecular clone (IMC) of pREJO.c/2864 (REJO, ARP-11746) was obtained through the NIH HIV Reagent Program, Division of AIDS, NIAID, NIH, contributed by Dr. John Kappes and Dr. Christina Ochsenbauer (55). REJO is a tier 2, clade B, T/F isolate. IMCs of SF162 (chronic; clade B, chronic, tier 1B) and CMU06 (acute; clade AE, acute, tier 2) were generated by cloning the Env into a pNL4.3 backbone to construct pNL-CMU06 and pNL-SF162, respectively (56). The human CD4 expression plasmid was kindly provided by Prof. F. Kirchhoff (57).

### Antibodies and lectins

The following antibody reagents used in this study were obtained through the NIH AIDS Reagent Program, Division of AIDS, NIAID, NIH: Anti-HIV-1 gp120 monoclonal PG9, PG16, PGT145, PGT121, PGT128, from IAVI (58); anti-HIV-1 gp120 monoclonal CH59 from Drs. Barton F. Haynes and Hua-Xin Liao (49); anti-HIV-1 gp120 monoclonal VRC01 from Dr. John Mascola (59); anti-HIV-1 gp120 Monoclonal b12 from Dr. Dennis Burton and Carlos Barbas (60); anti-HIV-1 gp120 monoclonal 3BNC117 from Dr. Michel C. Nussenzweig (61); anti-HIV-1 gp120 monoclonal 17b from Dr. James E. Robinson (62); anti-HIV-1 gp41-gp120 monoclonal 35O22, from Drs. Jinghe Huang and Mark Connors (63); anti-HIV-1 gp41-gp120 monoclonal PGT151 from Dr. Dennis Burton; anti-HIV-1 gp41 monoclonal 2F5 and 4E10 from Polymun Scientific (64). The V2i and V3 mAbs were obtained from the laboratory of Dr. Susan Zolla-Pazner (65–72). An irrelevant anti-anthrax mAb 3685 (73) was used as a negative control. The lectin Concanavalin A (ConA) was purchased from Vector Laboratories. The lectin Griffithsin was obtained through NIH HIV Reagent Program, Division of AIDS, NIAID, NIH, contributed by Drs. Barry O’Keefe and James McMahon.

### Virus production and purification

Infectious viruses were generated by transfecting 293T cells with pREJO, pNL-SF162 and pNL-CMU06 plasmids using jetPEI transfection reagent (Polyplus, New York, NY) (74). Supernatants were harvested after 48 hours and clarified by centrifugation and 0.45μm filtration. Virus infectivity was assessed on TZM.bl cells as described (56, 74). Briefly, serial two-fold dilutions of virus stock in 10% DMEM were incubated with TZM.bl cells (in duplicates for each dilution) in half-area 96-well plates in the presence of DEAE-dextran (12.5 μg/ml) for 48 hours at 37°C. Virus infectivity was measured by β-galactosidase activity (Promega, Madison, WI). Virus stocks were concentrated (20X) by ultracentrifugation over 20% (w/v) sucrose in 1X phosphate buffered saline (PBS) at 28,000 RPM for 2 hours in an SW-28 swinging bucket rotor (Sorvall, Thermofisher Scientific). Supernatants were decanted and pellets dried briefly before resuspension in PBS. Inactivation of virions was carried out using Aldrithiol-2 (AT-2) (21, 75). Briefly, 125 ul of sucrose-purified virus was incubated with 0.5 mM AT-2 in DMSO for 2 hours at 37^°^C, followed by centrifugation at 13,000 rpm for 2 hours. The supernatant was discarded, and the pellet re-suspended in 125 ul PBS. Inactivation was confirmed by measuring infectivity in TZM.bl cells and Env content was checked by Western blotting.

### Western blotting

To quantify and monitor the expression of Env in each virus preparation Western blot analyses were performed. The sucrose-purified virus particles were lysed, resolved by SDS-PAGE on 4–20% tris-glycine gels (Bio-Rad, Hercules, CA), and blotted onto membranes, which were then probed with antibodies. A cocktail of anti-human anti-gp120 MAbs (anti-V3: 391, 694, 2219, 2558; anti-C2: 841, 1006; anti-C5: 450, 670, 722; 1μg/ml each) was used to detect Env. MAb 91-5D (1μg/ml) was used to detect Gag p24. Membranes were developed with Clarity Western ECL Substrate (Bio-Rad, Hercules, CA) and imaged by a ChemiDoc Imaging System (Bio-Rad, Hercules, CA). Purified recombinant gp120 and p24 proteins were also loaded at a known concentration as controls and quantification standards. Band intensities were quantified using the Image Lab Software Version 5.0 (Bio-Rad, Hercules, CA).

### Coupling of fluorescent beads to virions

The coupling strategy used in this study is a standard method widely used in the Luminex based multiplex immunoassays and is published previously (76–79). We and others have used this protocol (xMAP Technology, Luminex Corp) and carboxylated beads to assess the Fc-mediated ADCP function of anti-HIV-1 and anti-COVID-19 antibodies (80–82). Here we applied this approach for coupling the virus to the beads. Sucrose-purified, inactivated virions were covalently coupled to carboxylate-modified microspheres (1.0 µm) using a two-step carbodiimide reaction with the xMAP Ab Coupling (AbC) Kit according to manufacturers’ instructions (Luminex, Austin, TX). Carboxylated beads purchased from Thermo Fisher (cat# F8823), were coupled to 125 μl of 20X concentrated virus preparations (∼36.4×10^9^ beads per reaction). Briefly, the stock microspheres were vortexed and sonicated to resuspend and disperse the microspheres and 12 μl was transferred to a tube containing 1200 ul of 1% BSA/1X PBS (per virus). The microspheres were washed twice with 500 μl of activation buffer followed by vortexing and sonication after each step. The microspheres were activated with 250 μL of activation buffer, 50 μL of 50 mg/ml Sulfo-NHS (N-hydroxysulfosuccinimide), 50 μL of 40 mg/mL ethyl dimethylaminopropyl carbodiimide hydrochloride (EDC) and incubated for 20 min at room temperature with end-to-end rotation. The microspheres were washed three times in activation buffer, then incubated with AT-2 inactivated virus in activation buffer for 2 hours at room temperature. We typically used a volume of 20X concentrated virus that equals to ∼175 ng total. The microspheres were subsequently washed and resuspended in 1.2 mL of 0.1% BSA/PBS and stored at 4°C until ready to use.

### Virus-associated Env binding assay (VAEBA)

The coupled microspheres were dispensed into 96-well plate (10 µl/well) and blocked with 100 ul of 3% BSA for 30 mins at 4°C. Plates were centrifuged at 2000 x g for 10 minutes and the supernatant was discarded. The microspheres were incubated with serially diluted mAbs for 30 min at 37°C followed by addition of anti-human-biotin (1:500 in 0.1% BSA/PBS) and incubated in the dark for 30 mins. Plates were washed 3 times with PBS and incubated with Streptavidin-PE (1:1000 in 0.1% BSA/PBS) for 30 min followed by resuspension in 200µl of 0.5% paraformaldehyde. Plates were washed 3 times with PBS after each step. Analysis was done with Attune NxT flow cytometer, and ≥30,000 events were collected in the phycoerythrin (PE)+ gate. Data analysis was carried out using FCS-Express software as follows: FITC positive microspheres were selected from a plot of forward-area vs. FITC (FSC-A/FITC-A) from which doublets were excluded in a forward scatter height vs forward scatter area plot (FSC-H/FSC-A). Geometric mean fluorescent intensity (MFI) values of PE+ beads representing anti-Env-stained virus particles coupled to FITC microspheres, were determined and plotted. Background MFI, as determined from microspheres stained without primary antibodies was subtracted from all Env-mAb pairs. Area under the titration curves was also calculated for select data sets. Unless otherwise noted in the figure legends the mAbs were tested at the following concentrations: 697, 830A, 2158, 3685 at 100 μg/ml; 2219, 2557 at 50 μg/ml; PG9, PG16, PGT145, CH59, VRC01, 3BNC117 at 25 μg/ml; 4E10 at 20 μg/ml; PGT151 at 10 μg/ml; NIH45-46 at 5 μg/ml.

For time-dependent binding studies, the assay was performed as above using biotinylated mAbs that were incubated with the virus-coupled beads for different times (0, 15, 30, 45, 60 and 75 min) followed by Streptavidin-PE. Data was acquired by Attune NxT flow cytometer, as above.

For conformational epitope exposure in response to mAbs, virus coupled microspheres were pre-incubated with titrated mAbs for 30 min at 37°C. Plates were centrifuged and further incubated with biotinylated mAbs (PG9 or 2219) for 30 min at 37°C followed by Streptavidin-PE. Data was acquired by Attune NxT flow cytometer, as above.

For virus-binding assays in the presence of 293T cells +/- human CD4, we transfected 293T cells with plasmids expressing full-length functional human CD4 (57), as below. Cells were harvested 24 hours post-transfection using trypsin-free cell-dissociation buffer, stained with Live/Dead fixable Aqua stain and labelled with Vybrant DiD cell labelling solution (Thermo Fisher Scientific) for 30 minutes at 4^°^C. Cells were distributed in 96 well V-bottom plates followed by addition of virus-coupled microspheres and serially diluted antibodies, and incubated for 30 minutes at 4^°^C. Binding was detected using biotinylated anti-human IgG and Streptavidin-PE. Data was acquired by Attune NxT flow cytometer, as above. Analysis was carried out using FCS-Express software using two gating strategies as follows: 293T cells and virus-coupled microspheres were selected from a plot of forward-area vs. side scatter-area (FSC-A/SSC-A). Gating strategy 1: DiD labelled cells were gated for live cells by selecting for Aqua-negative gating, followed by gating on cells that are positive for FITC (virus-coupled FITC microspheres) and geometric mean fluorescent intensity (MFI) of PE+ events, representing anti-Env-stained beads, were quantified. Gating strategy 2: FITC positive virus-coupled microspheres were gated for DiD positive cells, live cells were selected by Aqua-negative gating, and geometric mean fluorescent intensity (MFI) of PE+ beads/cells, representing anti-Env-stained beads, were quantified.

For conformational epitope exposure in response to lectin ConA, virus coupled microspheres were pre-incubated with titrated amounts of ConA for 30 min at 37°C. Plates were centrifuged and further incubated with mAbs (830A, PG9, 2219, NIH 45-46, VRC01, and negative control 3685) at one dilution for 30 min at 37°C followed by Streptavidin-PE. Data was acquired by Attune NxT flow cytometer, as above.

### Cell-associated Env binding assay

An assay to detect antibody binding to cell surface expressed Env was performed as described (56, 83, 84). Briefly, monolayers of 293T cells (4 × 10^6^) seeded in 100-mm tissue culture dishes were transfected with 20 μg of gp160 expression plasmid using jetPEI (DNA:jetPEI ratio of 1:3) following manufacturer’s instructions (Polyplus, New York, NY). Transfected cells were incubated for 24 hours at 37°C, washed with PBS, detached with trypsin-free cell-dissociation buffer, and resuspended in PBS containing 2% BSA. Cells were stained with Live/Dead fixable Aqua stain and distributed into 96-well V-bottom tissue culture plates (5×10^4^/well) for individual staining reactions. Cells were incubated with mAbs at concentrations detailed in figure legends. For detection of mAb binding, biotinylated goat anti-Human IgG Fc (1:500) followed by Streptavidin-PE (1:1000) was used. The cells were washed 3X with PBS-B (PBS plus 1% BSA) after each step and all incubation steps were performed on ice for 30 min. Cells were analyzed with a BD Fortessa flow cytometer, and >30,000 events were collected in the PE+ gate. Analysis was carried out using FCS-Express software as follows: 293T cells were selected from a plot of forward-area vs. side scatter-area (FSC-A/SSC-A) from which doublets were excluded in a forward scatter height vs forward scatter area plot (FSC-H/FSC-A). Live cells were selected by Aqua-negative gating, and geometric mean fluorescent intensity (MFI) of PE+ cells, representing anti-Env-stained cells, was quantified. Background MFI, as determined from cells stained without primary antibodies, was subtracted from all Env-mAb pairs.

### Lectin Capture Enzyme-Linked ImmunoSorbent Assay (ELISA)

The relative binding of mAbs to Env on virions was also measured by a lectin capture ELISA. Briefly, half-area high-binding ELISA plates (Costar) were coated with five-fold titrated amounts of ConA lectin (50 – 0.08 μg/ml in PBS) for 1 hour at 37^°^C. Sucrose pelleted REJO virus was AT-2 inactivated as above and added to the ConA coated plate. The plate was incubated for 1 hour at 37^°^C, washed, and blocked with 5% non-fat dry milk in whey solution for 1 hour at room temperature or overnight at 4°C. Antibodies were incubated with captured virus at a single dilution (830A, 2219, PG9, 3685 at 50 μg/ml; VRC01 at 25 μg/ml and NIH45-46 at 2.5 μg/ml), followed by biotinylated anti-human IgG and Streptavidin-HRP. The bound mAbs were detected with SuperSignal ELISA Pico Chemiluminescent Substrate (Thermo Fisher Scientific) and relative luminescence units (RLUs) were measured using a BioTek Cytation luminometer. For the lectin Griffithsin (GRFT), plates were coated with GRFT at 50 μg/ml, incubated with virus particles, and probed with five-fold serially diluted mAbs (830A, 2219, PG9, 3685 at 50 μg/ml; VRC01 at 25 μg/ml and NIH45-46 at 2.5 μg/ml). The bound mAbs were detected as above. The plates were washed 3X with PBS-T (PBS plus 0.05% Tween 20) after each step.

### Neutralization assay

Virus neutralization was measured using a β-galactosidase-based assay (Promega) with TZM.bl target cells (85). Serial dilutions of mAbs were added to the virus in half-area 96-well plates (Costar, Corning, NY) and incubated for designated time periods (0 min and 60 min) at 37°C. TZM.bl cells were then added along with DEAE-dextran (12.5 μg/ml; Sigma). After a 48-hour incubation for each assay condition, at 37°C and in a 5% CO2 incubator, the β-galactosidase activity was measured. Each condition was tested in duplicate. Assay controls included replicate wells of TZM.bl cells alone (cell control) and TZM.bl cells with virus alone (virus control). The highest antibody concentrations tested were based on known neutralization titers and varied per mAb. PG9 and 2219 were analyzed in the range of 50 μg/ml to 0.78 μg/ml, PGT145 was analyzed in the range of 5 μg/ml to 0.001 μg/ml, and NIH45-46 was analyzed in the range of 2.5 μg/ml to 0.004 μg/ml, while 830A and 3685 were tested in the range of 100 μg/ml to 0.006 μg/ml. Percent neutralization was determined on the basis of virus control under the specific assay conditions. The virus input corresponded to titrated stocks yielding 150,000 to 200,000 relative luminescence units (RLU).

### Statistical analysis

All statistical analyses were performed with GraphPad Prism 9.4.1 (GraphPad, San Diego, CA). ANOVA and Mann-Whitney t-tests were performed as appropriate and are mentioned in the figure legends.

## Results

### Detection of REJO Env on HIV-1 particles

To analyze the representation of Env epitopes on virus particles, we established a flow cytometry-based assay. The assay developed is based on (a) coupling the virions to fluorescent microspheres; (b) staining virions immobilized on microspheres with antibodies targeting different epitopes; (c) staining the resulting complex with biotinylated anti-human IgG and streptavidin-PE; and (d) detecting Env-Ab interaction via flow cytometry (Fig. 1A).

**Figure. 1.**
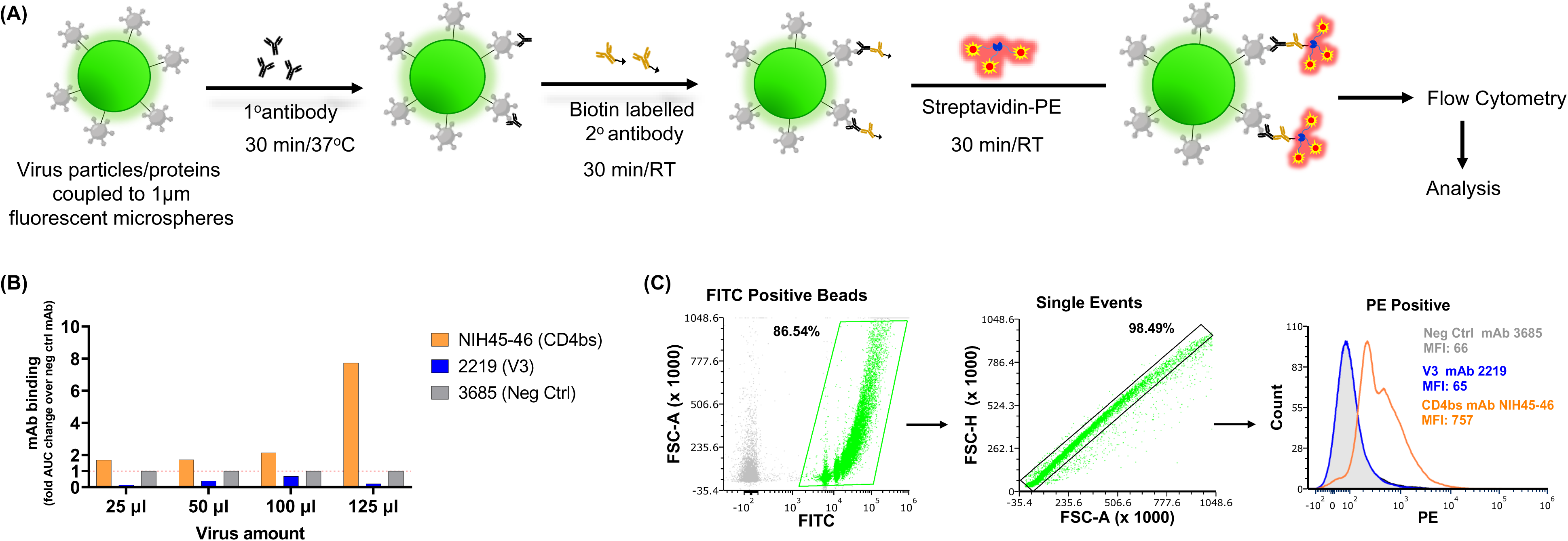
Virus-binding assay for detection of antibody binding to Env expressed on virions. (A) Schematic of the assay; (B) Optimization of virus amount to be coupled to the microspheres; (C) Representative dot plots and histogram showing the gating strategy for analysis of antibodies binding to virus particle by flow cytometry. The mAbs were tested at following concentrations: NIH45-46 at 5 μg/ml; 2219 and 3685 at 50 μg/ml. 1°, primary; 2° secondary.

Initially, we used REJO—a T/F, clade B, tier 2, HIV-1 isolate—and select monoclonal antibodies (mAbs) that recognize the conformational CD4bs (NIH45-46) and linear V3 (2219) epitopes on Env. A non-HIV-1 mAb 3685 was also used as a negative control. To optimize the amount of sucrose-pelleted virus required to couple with the microspheres, we tested four different amounts (25, 50, 100 and 125 μl corresponding to ∼35, 70, 142 and 177 ng total Env, respectively) of 20X concentrated virus preparations. The Env content was measured by Western blot (Fig. S1) as in (56, 74).

At 125 μl of virus, the binding of positive control mAb NIH45-46 was significantly enhanced while the negative control mAb 3685 background binding levels were low (Fig. 1B). Thus, based on the fold AUC change over the non-HIV-1 mAb 3685, we elected to use 125 μl of 20X concentrated virus preparation, equivalent to 177 ng Env. At this amount, the negative control mAb stained the virions minimally (1.7%) (Fig. 1C). The mAb NIH45-46 bound 84% of REJO virions, while only 1.8% of particles bound to V3 mAb 2219 (Fig. 1C). Next, we used the assay to analyze the Env epitopes present on the REJO virus particles using the extended mAb panel (Table 1) that covers most of the major Env domains.

**Table 1.**
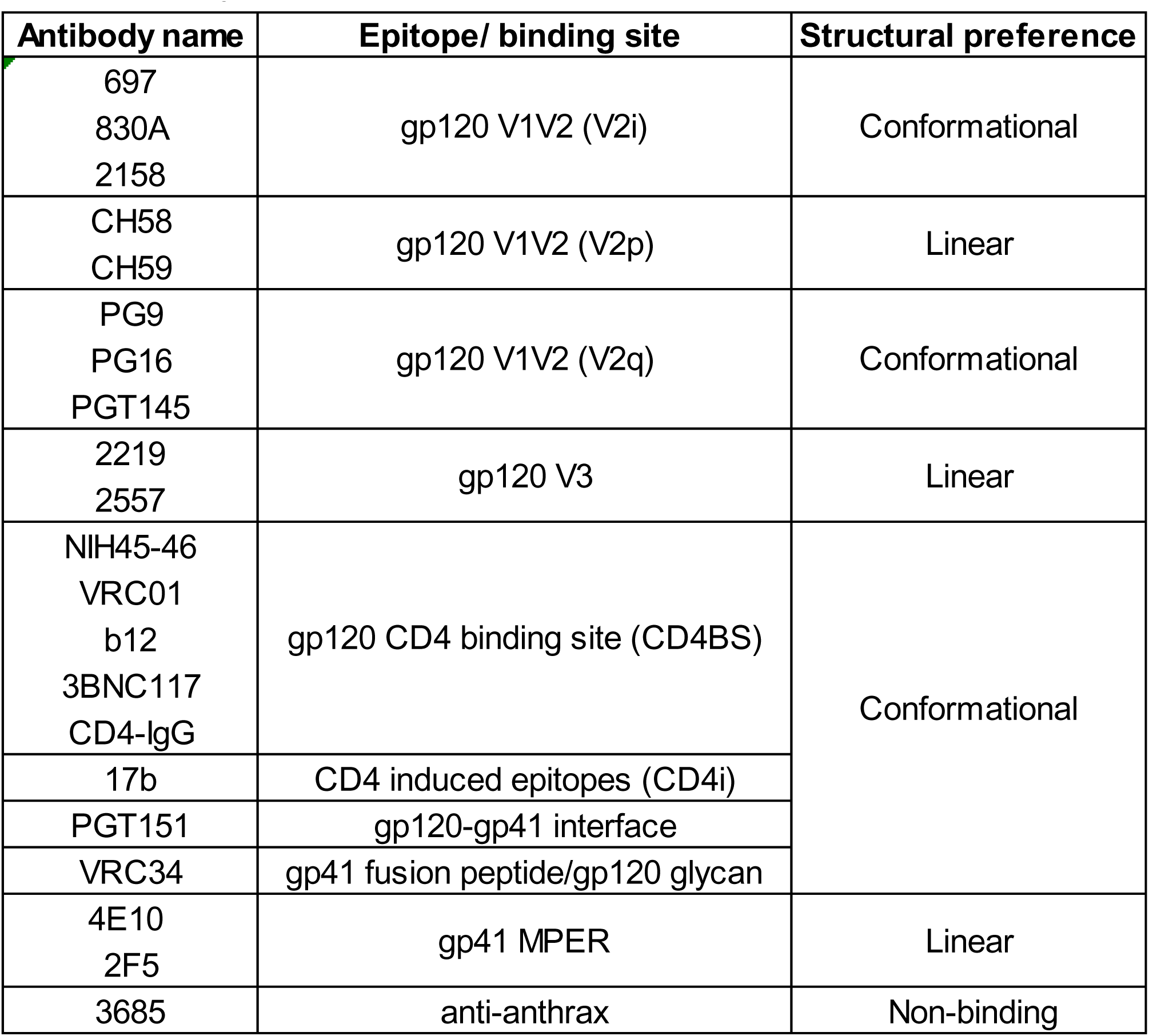
Binding specificity of the monoclonal antibodies (mAbs) used in the study.

| **Antibody name** | **Epitope/ binding site** | **Structural preference** |
| --- | --- | --- |
| 697<br>830A<br>2158 | gp120 V1V2 (V2i) | Conformational |
| CH58<br>CH59 | gp120 V1V2 (V2p) | Linear |
| PG9<br>PG16<br>PGT145 | gp120 V1V2 (V2q) | Conformational |
| 2219<br>2557 | gp120 V3 | Linear |
| NIH45-46<br>VRC01<br>b12<br>3BNC117<br>CD4-IgG | gp120 CD4 binding site (CD4BS) | Conformational |
| 17b | CD4 induced epitopes (CD4i) |  |
| PGT151 | gp120-gp41 interface |  |
| VRC34 | gp41 fusion peptide/gp120 glycan |  |
| 4E10<br>2F5 | gp41 MPER | Linear |
| 3685 | anti-anthrax | Non-binding |

The REJO virus-bound microspheres were treated with titrated amounts of mAbs and the assay was conducted as outlined in the Methods section. All three V2i mAbs (697, 830A, and 2158) bound to Env on virions, with mAb 830A exhibiting the strongest binding (Fig. 2A-C, S2). In contrast, binding of trimer-preferring V2q bNAbs (PG9, PG16, and PGT145) and peptide recognizing V2p mAbs (CH58 and CH59) was minimal and similar to the negative control mAb 3685 (Fig. 2A-C). Binding of V3 specific mAbs 2219 and 2557 also was similar to 3685 (neg control mAb) (Fig. 2). Recognition of CD4bs epitopes by most mAbs, except NIH45-46, was also negligible. The gp120-gp41 interface (PGT151) and gp41 MPER mAbs (4E10) displayed efficient binding. Higher binding of MPER specific 4E10 mAb may be either due to lipid cross-reactivity, increased exposure of the gp41 base, or presence of gp41 stumps (21, 86, 87). Notably, PGT151 which is known to be trimer-specific and binds only to properly formed, cleaved trimers, had comparable binding to the V2i mAb 830A (88).

**Figure 2.**
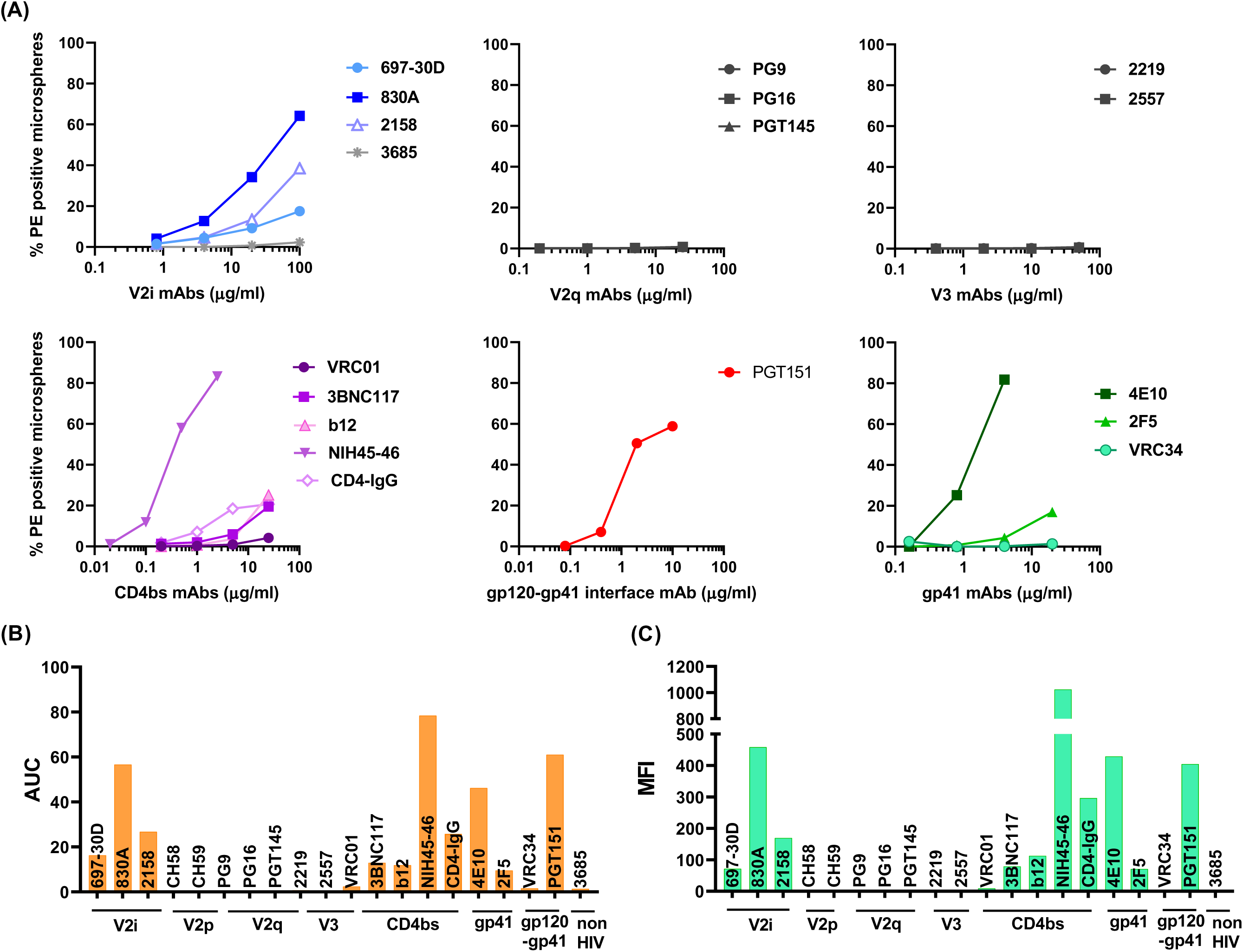
Binding of mAbs to REJO Env on virions. (A) Microspheres coupled to REJO virus were reacted with serially diluted mAbs targeting different Env epitopes. (B) Area under the curve (AUC) were calculated from the curves in A and plotted. (C) Geometric mean fluorescent intensity (MFI) of each mAb tested at a single dilution are plotted. Virus coupled microspheres stained with biotin and PE alone were used to set the background MFI and subtracted.

To compare the relative levels of each epitope exposed on virus particles, we calculated AUC values from the titration curves. Figure 2B represents the overall presentation of different epitopes on the virions as measured by mAb binding. A similar pattern was also observed when geometric mean fluorescent intensities (MFI) of mAbs tested at the highest dilution were plotted (Figure 2C).

### Pattern of Env epitope exposure on virions from different clades, tiers, or infection stage

We next tested two other HIV-1 isolates: SF162 (chronic; clade B, tier 1) and CMU06 (acute; clade AE, tier 2) using the same panel of mAbs. In case of SF162 virions, the V2i mAbs 697 and 830A showed greater levels of binding than 2158 (Fig 3, S3). As seen with REJO, V2q, V2p, and V3 mAbs showed negligible binding to SF162 virions and were similar to negative control mAb 3685 (Fig 3, S3). All CD4bs mAbs tested showed binding to SF162 virions that was above background level. In case of CMU06, a similar pattern was observed with mAbs 830A, PGT151, and 4E10 showing highest binding levels, while little to no binding was seen with the mAbs targeting the V2q and V3 epitopes (Fig 4, S4). Negligible binding was seen with the negative control mAb 3685. Overall, the binding strength of mAbs to the SF162 Env on virions was higher compared to CMU06 and REJO. The epitopes recognized by mAbs 830A, PGT151, and 4E10 were consistently accessible on all three viruses tested in this study. Differences in the recognition of the CD4bs mAbs among the three isolates were also evident - mAbs 3BNC117 and VRC01 showed better binding to SF162 than the other two viruses. The CD4bs mAb NIH45-46 is a more potent clonal variant of VRC01 with high sequence and structural similarities to VRC01, but has a distinct mode of binding to gp120 (89). The potency of the CD4bs mAbs tested here is in the order NIH45-46 > 3BNC117 > VRC01 (90). Differences in binding, in that order, are evident in the binding data shown in Fig 2-4.

**Figure 3.**
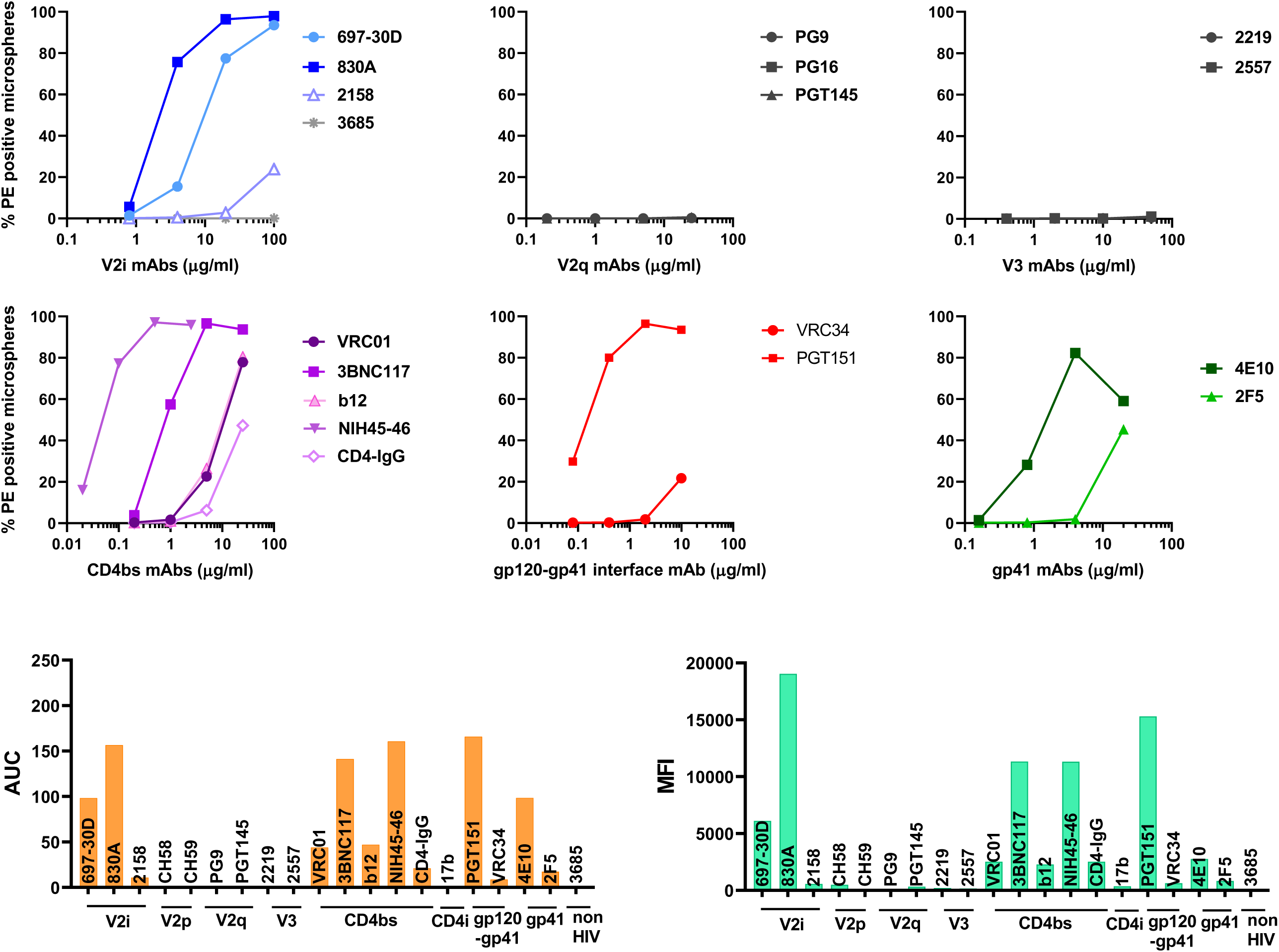
Binding of mAbs to SF162 Env on virions. (A) Microspheres coupled to SF162 virus were reacted with serially diluted mAbs targeting different Env epitopes. Virus coupled microspheres stained with biotin and PE alone were used to set the background MFI and subtracted. (B) Area under the curve (AUC) were calculated from the curves in A and plotted. (C) Geometric mean fluorescent intensity (MFI) of each mAb tested at a single dilution are plotted.

**Figure 4.**
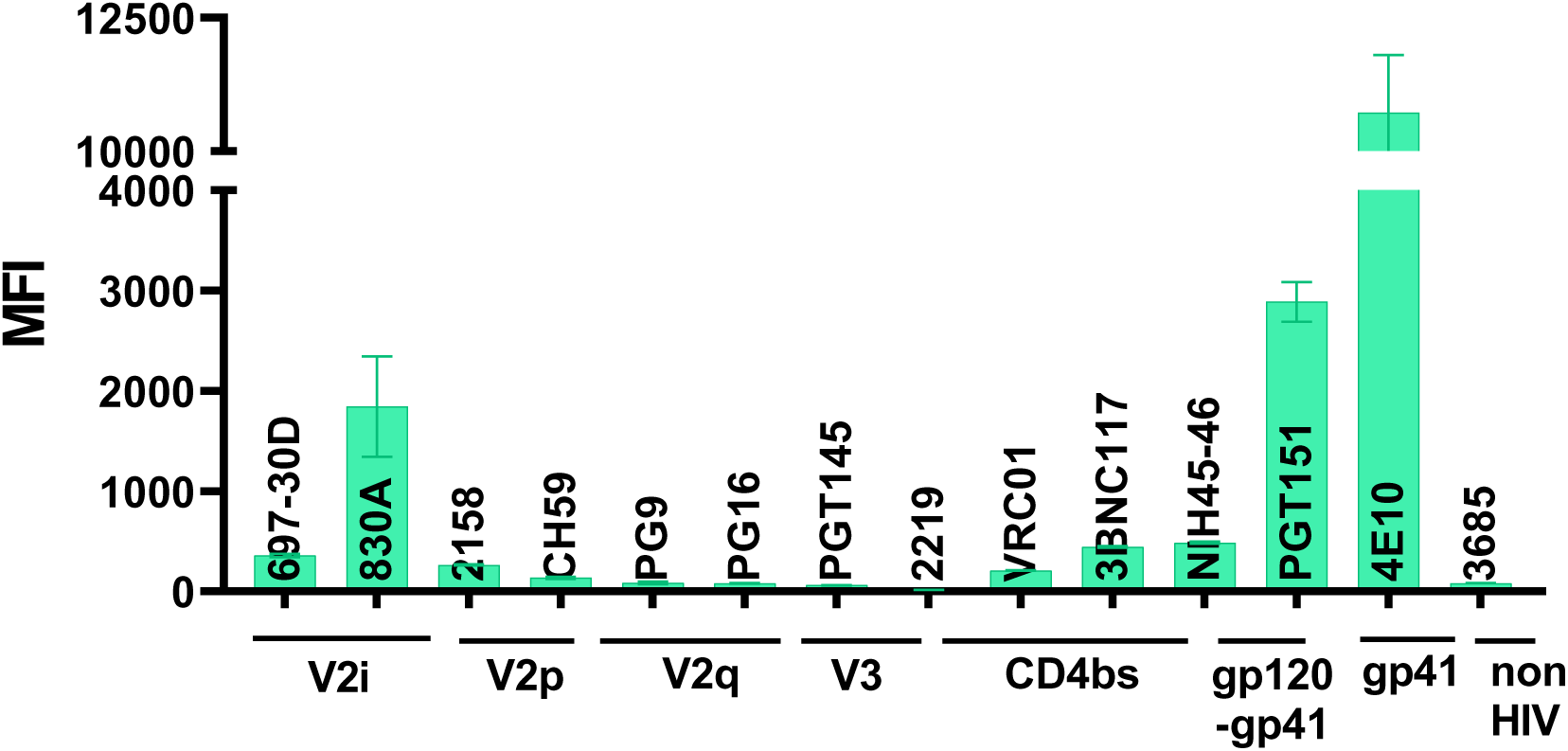
Binding of mAbs to CMU06 Env on virions. (A) Microspheres coupled to CMU06 virus were reacted with mAbs targeting different Env epitopes. Geometric mean fluorescent intensity (MFI) of each mAb tested at a single dilution are plotted. Virus coupled microspheres stained with biotin and PE alone were used to set the background MFI and subtracted.

### Binding profile of mAbs to Env expressed on the cell surface is different from Env on virions

We next sought to compare the binding of mAbs to virions with binding to transiently expressed Env on cells. We transfected 293T cells with gp160 expression plasmids and probed Env expressed on the cell surface with the mAbs in Table 1. A different profile was observed when staining of Env on virus particles was compared to Env expressed on transfected cells (Fig. 5, S5, S6, and S7). All mAbs tested bound to Env expressed on cells, while this was not the case with HIV-1 particles (Fig. 2-4). The V2q, V2p, and V3 mAbs which showed negligible binding to Env on virions displayed efficient binding to REJO and CMU06 Env expressed on 293T cell surface. Thus, Env recognized by V2q mAbs are efficiently expressed however, the mechanism of Env incorporation may affect its conformation and expression at the virion surface. In addition, Env on producer cells may also be populated by uncleaved Env monomers at the cell surface that are not incorporated into virions (91), which may also account for the observed differences in binding. The CD4bs mAbs also displayed greater binding to cell-surface expressed Env compared to Env on virus particles. Env from all three isolates also bound efficiently to PGT151, which binds only to properly formed, cleaved trimers (92). Thus, the binding profile of mAbs to Env differs based on the location of the Env (cell surface versus virus surface).

**Figure 5.**
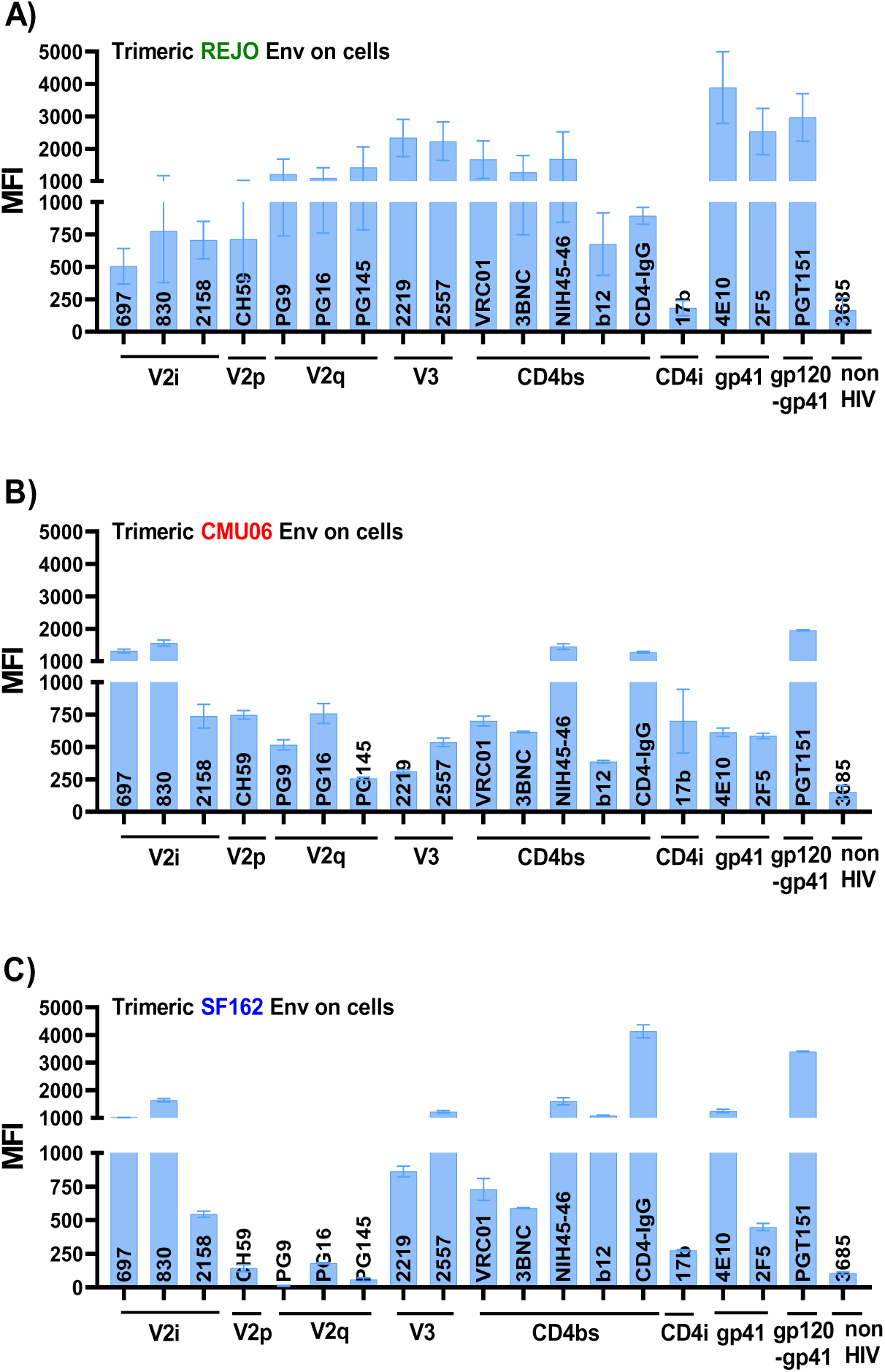
Binding of mAbs to trimeric Env expressed on cell surface. 293T cells were transiently transfected with plasmid encoding for full length gp160 of (A) REJO, (B) CMU06 and (C) SF162. Cells were analyzed for binding of different mAbs 24-hours post-transfection. Geometric mean fluorescence intensity (MFI) and SD from duplicates or triplicates in one experiment are shown. Transfected cells stained with biotin and PE alone were used to set the background MFI and subtracted. The mAbs were tested at following concentrations: 697, 830A, 2158, 3685 at 100 μg/ml; 2219, 2557 at 50 μg/ml; PG9, PG16, PGT145, PGDM1400, CH59, VRC01, 3BNC117, b12 at 25 μg/ml; 4E10, 2F5 at 20 μg/ml; PGT151 at 10 μg/ml; NIH45-46 at 5 μg/ml.

### Prolonging the mAb-virion interaction time does not unmask the occluded epitopes

Binding of antibodies to Env is essential in order for them to neutralize the virus. In a typical standard neutralization assay, virus and mAbs are pre-incubated for 1 hour at 37°C prior to adding the target cells. In our Env binding assay, mAbs-virus were incubated for only 30 min at 37°C, prompting us to test if increasing the mAb-virus interaction time would facilitate binding, especially for the mAbs (V2q, V3, and CD4bs) that showed little to no binding after 30 minutes.

To test this idea, microspheres coupled with REJO virus were incubated with select mAbs: V2i (830), V2q (PG9), V3 (2219), CD4bs (NIH45-46), or control mAb (3685) for various periods of time (0 to 75 min). The binding of mAbs to the Env on REJO virus was measured by flow cytometry (Fig. 6, S8). Little binding to REJO Env was observed for all mAbs when no incubation (0 min) was allowed, but a significant linear increase in binding of 830A and NIH45-46 mAbs was detected over time (Fig. 6, S8). The V2q mAb PG9 and V3 mAb 2219 showed no increase in binding to REJO Env at any time point, indicating that epitopes of these mAbs remained occluded on the virus even after 75 min of incubation (Fig. 6). As expected, no binding was detected for the negative control mAb 3685 at any time point.

**Figure 6.**
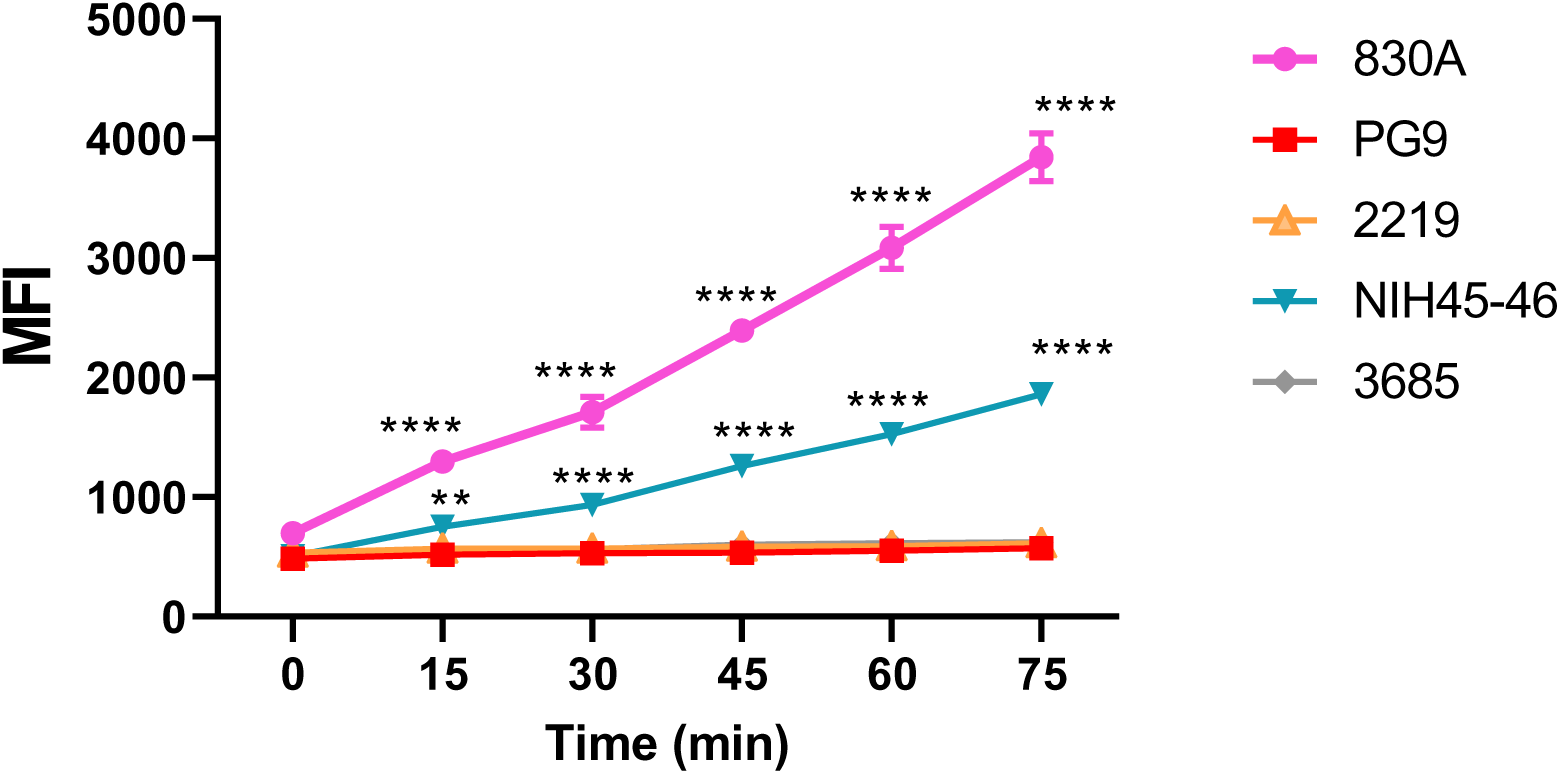
Time-dependent increase of Env binding. Microspheres coupled to REJO virus particles were treated with mAbs at 37°C for various time from 0 to 75 min. Binding was detected as in Figure 2 and geometric mean fluorescence intensity (MFI) are shown. Virus coupled microspheres stained with biotin and PE alone were used to set the background MFI and subtracted. The mAbs were tested at following concentrations: 830A, 3685 at 100 μg/ml; 2219 at 50 μg/ml; PG9 at 25 μg/ml; NIH45-46 at 2.5 μg/ml. ****, p= <0.0001; **, p= 0.0026 vs t = 0 min by ANOVA.

### Masked Env epitopes can be exposed by pre-binding of other mAbs

We wondered if microsphere-virion coupling method might be contributing to the inability to detect binding of mAbs to these epitopes. If that stands correct then binding of V2q mAb PG9 or V3 mAb 2219 will remain low on virions even under conditions that are known to conformationally unmask these epitopes (93, 94). We tested this idea by pre-treating the virus coupled microspheres with select mAbs followed by detecting the binding of V2q (PG9) and V3 (2219) mAbs. We incubated the microsphere-coated virions first with mAbs 697, 830A, 2158, 447, or 2219 to induce structural changes in Env. After washing away the unbound mAb, we probed for V2q epitope exposure using biotinylated PG9 mAb. Non-biotinylated PG9 and non-HIV-1 mAb 3685 were used as controls. As seen in Fig. 7 and S9, binding of all three V2i mAbs (697, 830A, and 2158) presumably induced an allosteric conformational change that exposed the PG9 binding epitopes. This concurs with the published study showing that V2i 830A and 2158 increase the binding of PG9 on A244 gp120 (94). In contrast, the V3 mAbs (447 and 2219) did not alter PG9 binding and were comparable to the negative control mAb 3685. Since PG9 epitopes are also occluded on virions, negligible binding was seen in this control as well. Thus, these data further support the notion that PG9 epitopes are conformationally masked on virions. We next evaluated if V3 epitopes can be similarly exposed. The V3 loops are tucked beneath V1V2, thus Abs that induce any movement in the V1V2 loops should expose the V3 epitopes. We made use of the following CD4bs mAbs: the bNAb NIH45-46 that binds strongly to Env on virions, the non-neutralizing mAb 654 that is known to expose V3 loops, and the V2i mAb 830A (93, 94). Non-biotinylated 2219 and 3685 were used as controls. Binding of 830A and 654 exposed the V3 epitopes, allowing the binding of biotinylated 2219 mAb. In contrast, NIH45-46, despite binding well to REJO virions, did not induce changes in Env that might expose the 2219 epitopes (Fig 7, S9). The 2219 epitopes are not available on virions in the absence of allosteric alterations; thus, no binding was detected in this control.

**Figure 7.**
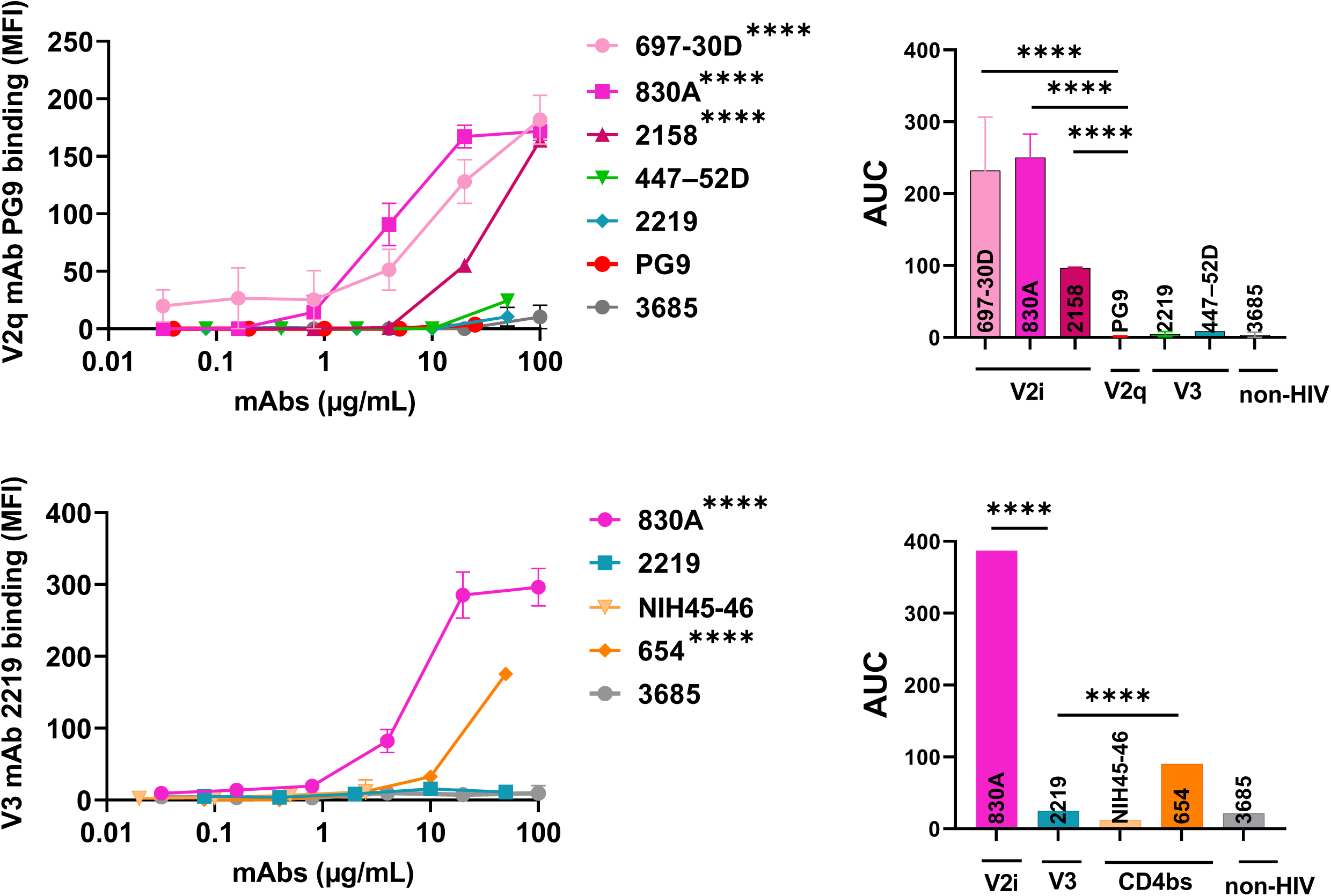
Changes in reactivity of PG9 and 2219 mAbs. Microspheres coupled with REJO virus were incubated with titrated amounts of antibodies followed by biotinylated V2q mAb PG9 (25 μg/ml) or biotinylated V3 mAb 2219 (50 μg/ml). Binding was detected with streptavidin-PE and geometric mean fluorescent intensity (MFI) are plotted. Virus coupled microspheres stained with biotin and PE alone were used to set the background MFI and subtracted. **** p< 0.0001 by ANOVA.

### Neutralization of REJO and CMU06 viruses by bNAbs

The findings from above suggest that allosteric changes, but not time, can allow exposure of the epitopes targeted by the V2q mAb PG9. However, given the reported neutralization breadth and potency of PG9, the little to no binding of PG9 to the Env on virion surface was perplexing (58, 59). This led us to speculate that perhaps the interaction of virus with the target cells induces similar changes in Env that allows bNAbs such as PG9 to bind to their epitopes. In such a case, we would expect that neutralization of HIV-1 with PG9 should be similar in the presence or absence of a virus-mAb pre-incubation step.

Indeed, as seen in Fig. 8, V2q bNAbs PG9 and PGT145 were able to neutralize REJO virus with equal efficacy independent of a pre-incubation step. Among the CD4bs mAbs, neutralization by 3BNC117 and NIH45-46 was also similar at both t = 0 and 60 minutes, while VRC01 had slightly better neutralization at t =60 min (AUC 111) compared to t = 0 (AUC 85); however, the difference was not statistically significant. These data suggest that interaction of virus with target cells appears to induce localized antigenic changes in Env on virions, allowing for bNAbs to latch onto their epitopes. This is the first time, to the best of our knowledge that the V2q and CD4bs mAbs tested here are shown to require pre-interaction of virus with target cells to exhibit their neutralization effect. Interestingly, mAbs such as 830A and 4E10 show strong binding to their respective epitopes on virions. However, these mAbs fail to neutralize the REJO virus. One possible explanation for this observation may be that the 830A and 4E10 epitopes that are available for binding on virions are associated with non-functional Env. Similar results were also observed when neutralization of CMU06 virus was compared at t = 0 vs 60 min (Fig 9). Collectively, these data highlight the importance of gaining a better understanding of the multiple mechanisms that are utilized by HIV-1 to avoid interaction with host-mounted Abs and maintain infectivity. Also, these findings encourage the idea to re-evaluate the assay formats that are widely used in the field.

**Figure 8.**
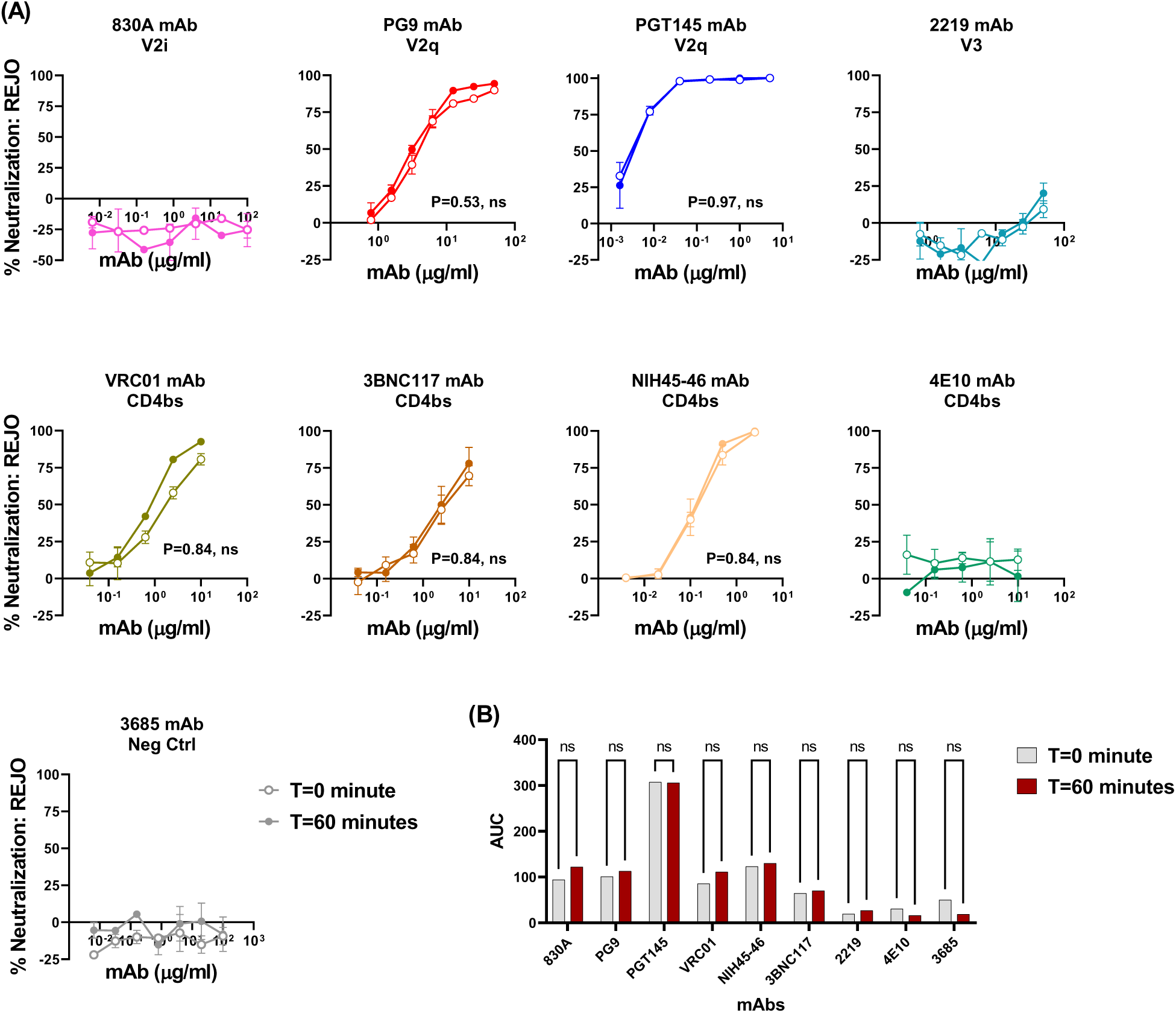
Neutralization of REJO virus. Virus neutralization was measured after REJO virus was incubated with serially diluted mAbs for the designated period of time at 37°C prior to the addition of TZM-bl target cells. Virus infectivity was assessed 48 h later based on β-galactosidase activity. (A) Neutralization curves are shown. Means and standard errors calculated from two different experiments (each in duplicate) are shown. Statistical analyses were performed on the neutralization curves reaching ≥50%. Comparison is made between neutralization curves at T = 0 vs T = 60 minutes. P= ns, not significant by nonparametric Mann-Whitney t-test. (B) Area under the neutralization curves were calculated and plotted as bar graph. Statistical analysis was performed on AUC graph by ANOVA (P = 0.99; ns, not significant).

**Figure 9.**
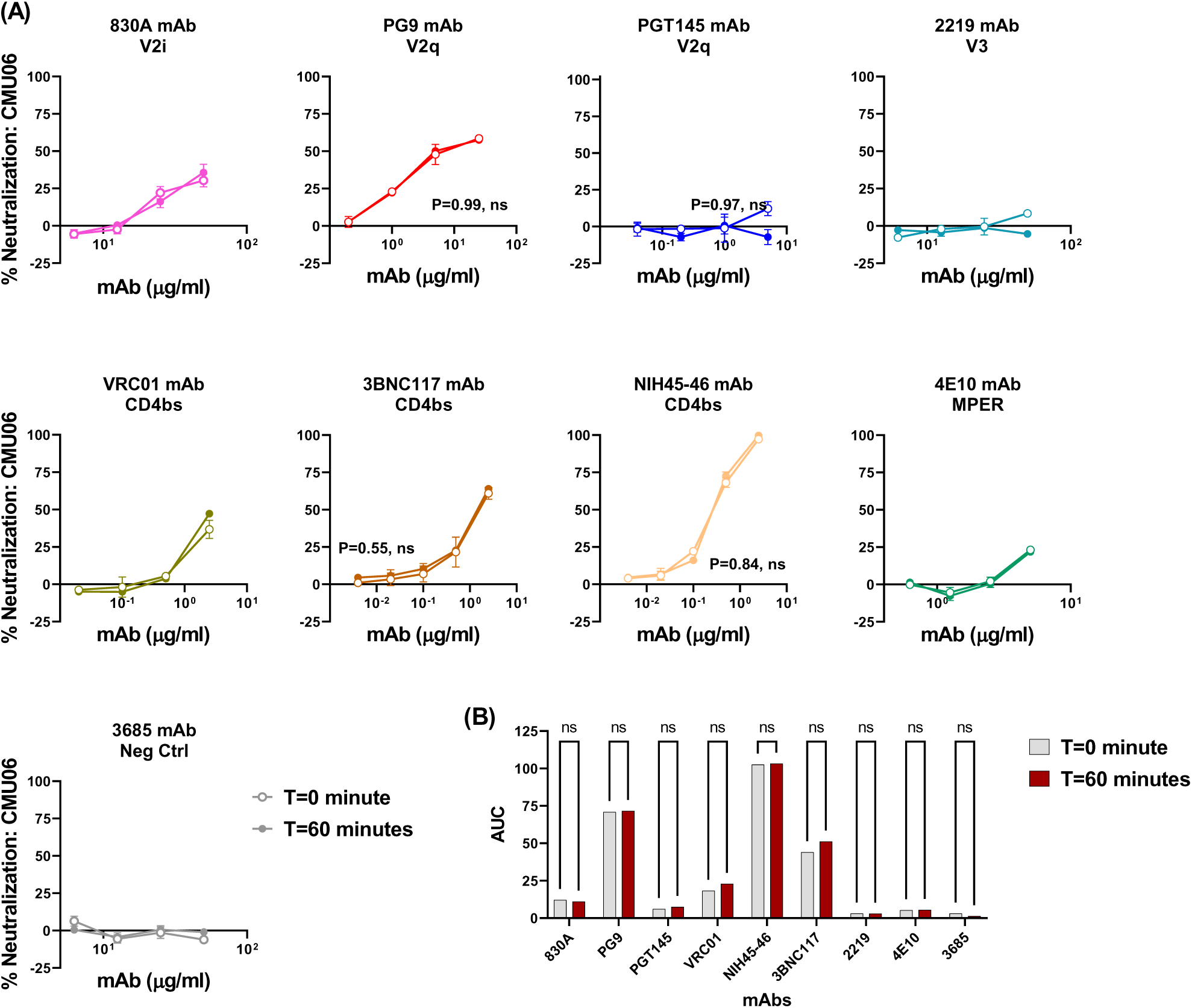
Neutralization of CMU06 virus. Virus neutralization was measured after CMU06 virus was incubated with serially diluted mAbs for the designated period of time at 37°C prior to the addition of TZM-bl target cells. Virus infectivity was assessed 48 h later based on β-galactosidase activity. (A) Neutralization curves are shown. Means and standard errors calculated from two different experiments (each in duplicate) are shown. Statistical analyses were performed on the neutralization curves reaching ≥50%. Comparison is made between neutralization curves at T = 0 vs T = 60 minutes. P= ns, not significant by nonparametric Mann-Whitney t-test. (B) Area under the neutralization curves were calculated and plotted as bar graph. Statistical analysis was performed on AUC graph by ANOVA (P = 0.99; ns, not significant).

### Influence of virus-cell interaction on Env epitope exposure

To further verify that virus interaction with host cells can indeed improve the exposure of Env epitopes on virions, we measured the binding of select mAbs to virion coupled fluorescent microspheres in the presence of host cells. The TZM.bl cells used for neutralization assays are HIV-1 permissive reporter HeLa cells modified to express the receptor (human CD4) and co-receptors (human CCR5 and CXCR4), allowing the virus to initiate infection in these cells (85). In addition, the surface of every eukaryotic cell is known to be covered by a dense and diverse array of cell surface glycans, the “glycocalyx” (95). Studies have shown that host cell glycocalyx can capture HIV-1 via specific, non-electrostatic oligomannose-GlcNAc glycan-glycan interactions to support viral entry (96, 97). To test the idea that host-cell interaction exposes Env epitopes, we used 293T cells that are non-permissive to HIV-1 entry for this assay. We transfected 293T cells with a plasmid expressing human CD4 (57, 98), to study the effect of surface CD4 on Env epitope exposure. In addition, we used un-transfected 293T cells, to study how glycan-glycan interaction (i.e, host glycocalyx and Env glycan interactions) may affect epitope exposure on virions. As seen in Fig 10 and S10, differences in binding of mAbs are evident, particularly for PG9 and 2219 that is relatively higher in the presence of 293T cells, compared to 293T-hCD4 cells.

In our previously published study, we have shown that lectin interaction with HIV-1 Env glycans can influence the exposure of V2i and V3 epitopes (99). To test if lectins can also expose the V2q epitopes on virions, we used Concanavalin A (ConA), which is a carbohydrate-binding protein extracted from jack-bean (*Canavalia ensiformis*) seeds and binds to the mannose residues of various glycoproteins (100, 101). ConA binds specifically to the α-mannose/α-glucose moieties on N-linked oligomannose and complex-type glycans and is not known to bind O-glycans on glycoproteins (102). We incubated the microsphere-coupled virions first with titrated amounts of ConA followed by select mAbs (830A, PG9, 2219, NIH45-46, VRC01, and 3685) at a single concentration. As seen in Fig 11 and S11, binding of all anti-HIV-1 mAbs including PG9, VRC01, and 2219 was detected in the presence of ConA. We further verified these findings using an alternative ELISA-based approach where we do not couple the virus particles to microspheres. We used ConA to capture the virus particles and detected the binding of different mAbs to captured virions by ELISA. As seen in Fig 12A, virions captured by ConA also displayed improved exposure of the PG9 and VRC01 epitopes, similar to the pattern observed in the flow cytometry based virus-binding assay (Fig 11). In contrast, another lectin, Griffithsin (GRFT), which is specific for α1–2 mannose on oligomannose glycans, exposed 2219 and VRC01 but not PG9 epitopes (fig 12B). Thus, these data further support the notion that HIV-1 masks the vulnerable epitopes on its Env and different factors (e.g., glycan-glycan and lectin-glycan interactions) may modulate the antibody accessibility of these epitopes on the virus particles.

**Figure 10.**
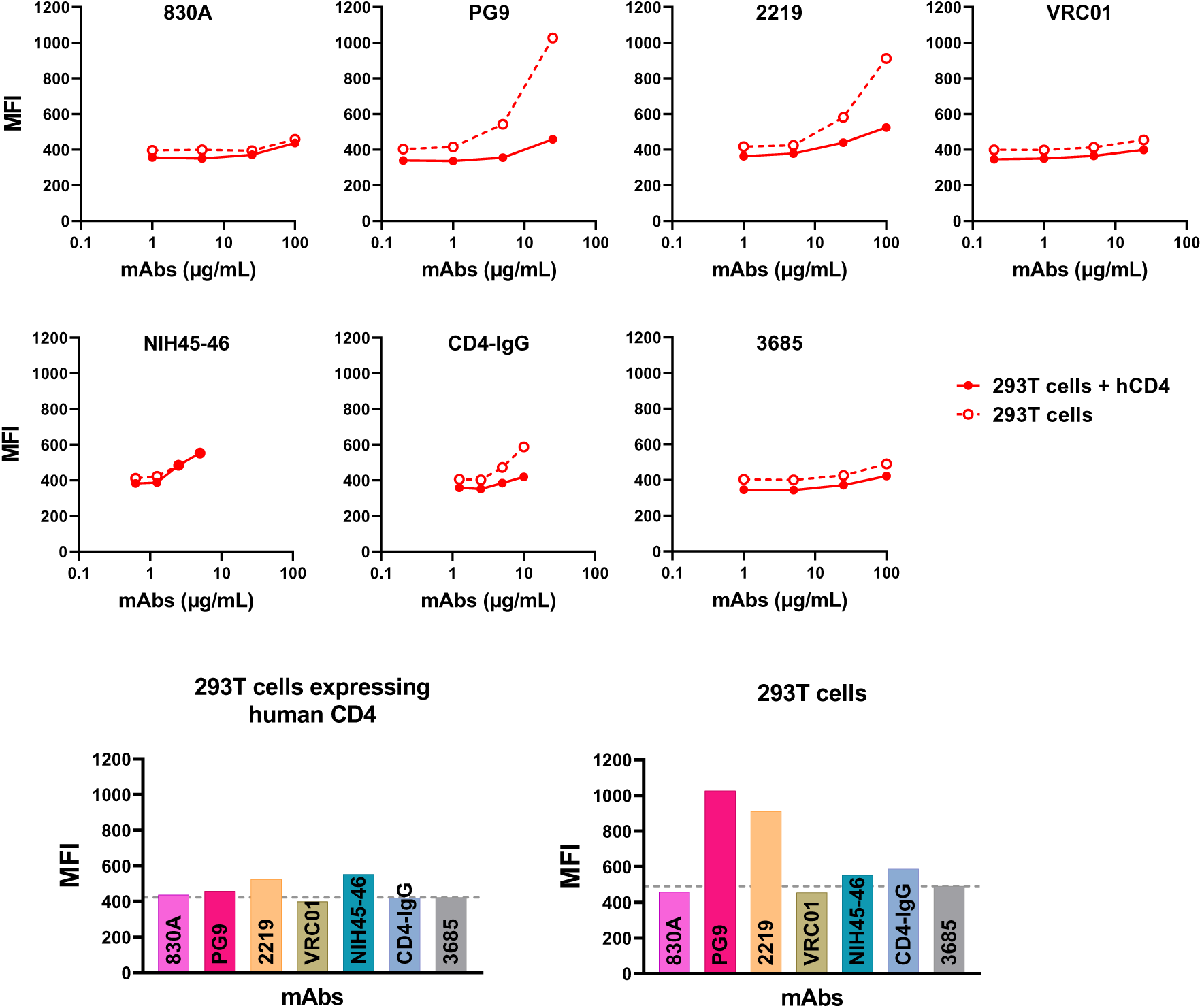
Influence of virus-cell interaction on Env epitope exposure. Binding of mAbs to virus-coupled microspheres were measured in the presence of 293T cells by flow cytometry. Cells were transfected or not transfected with human CD4 plasmid for 24 h and treated with virus-coupled microspheres and V2i, V3, V2q, CD4bs or control mAbs at 4°C. MAb binding was detected with biotin-conjugated anti-human IgG and PE-Streptavidin. Percent PE positive microspheres from one experiment are shown. Gating strategy used for data analysis is shown in Fig S10. MAb titration curves and bar graphs showing geometric mean fluorescent intensity (MFI) of each mAb tested at a highest dilution are plotted. Gray dotted line represents MFI of negative control mAb 3685.

**Figure 11.**
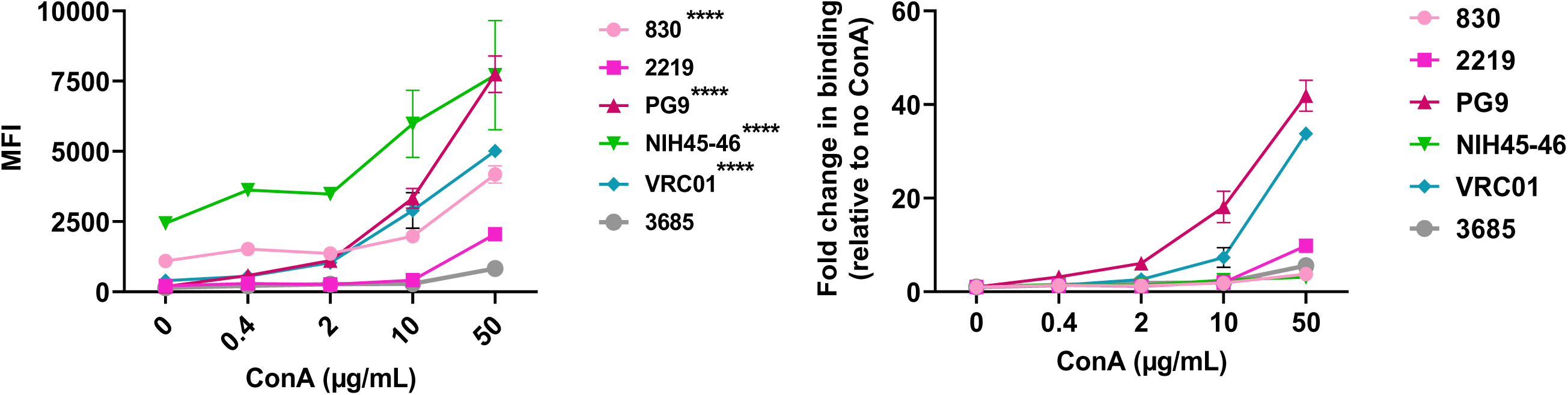
Effect of lectin Concanavalin A on exposure of Env epitopes. Microspheres coupled with REJO virus were incubated with five-fold titrated amounts of lectin Concanavalin A (ConA) (50 - 0.08 μg/ml) followed by mAbs at single dilutions (830A, 2219, PG9, 3685 at 50 μg/ml; VRC01 at 25 μg/ml and NIH45-46 at 2.5 μg/ml). (50 μg/ml). Binding was detected with streptavidin-PE. Geometric mean fluorescent intensity (MFI) and fold change in binding relative to no ConA are shown. Virus coupled microspheres stained with biotin and PE alone were used to set the background MFI and subtracted. **** p= 0.0002, * p= 0.039 by ANOVA relative to negative control mAb 3685.

**Figure 12.**
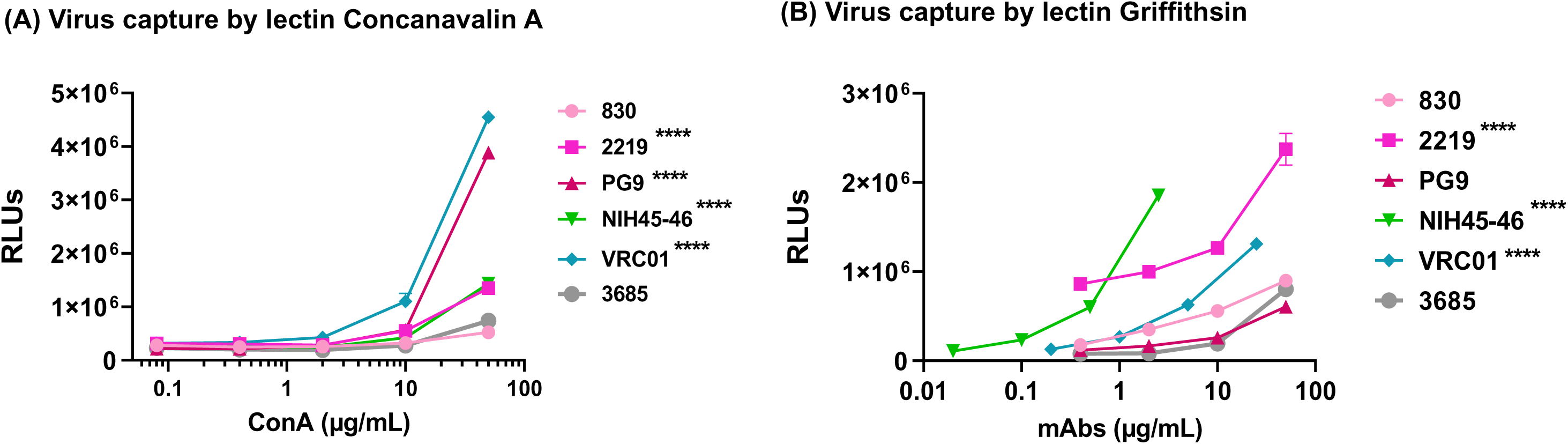
Effect of Lectins on Env epitope exposure in ELISA. Sucrose pelleted REJO virus was captured by (A) five-fold titrated lectin ConA (50 - 0.08 μg/ml) and reacted with different mAbs at one dilution (830A, 2219, PG9, 3685 at 50 μg/ml; VRC01 at 25 μg/ml and NIH45-46 at 2.5 μg/ml). (B) Sucrose pelleted REJO virus was captured by lectin Griffithsin (50 μg/ml) and reacted with five-fold titrated mAbs (830A, 2219, PG9, 3685 at 50 μg/ml; VRC01 at 25 μg/ml and NIH45-46 at 2.5 μg/ml). AUC values were calculated from binding curves. AUC changes relative to the respective WT are shown. Statistical analysis was performed on binding curves by 2-way ANOVA (**** p < 0.0001) relative to negative control mAb 3685.

## 4. Discussion

This study focused on investigating the antigenic landscape of HIV-1 Env on the infectious virus particles that are encountered by the host immune system. We developed a flow cytometry-based bead assay to quantitate the relative levels of Env epitopes that are exposed on HIV-1 particles. Although bead-based assays have been reported before, our approach varies from previous assays in that it alleviates the requirement of producing fluorescent virus particles (103). In addition, other assays make use of two anti-HIV-1 Abs, one Ab to capture the virus particles and other Ab to detect binding (104). As shown in this study, application of two Abs may skew the results and fail to differentiate between the actual mAb binding and binding due to the conformational changes in Env influenced by the capture Ab. Furthermore, the assay described in this study can be easily translated to study other viruses (e.g., SARS-CoV-2, Influenza etc.,) and antigens in different formats (e.g. gp120, gp140 etc).

Findings from this study show that epitopes targeted by mAbs like 830A (V2i), PGT151 (gp120-gp41 interface) and 4E10 (gp41) are present at relatively higher levels on the surface of REJO, CMU06 and SF162 viruses, suggesting that elements that regulate Env epitope abundance are likely to be conserved among isolates. In contrast, the V2q and V3 epitopes are inaccessible on the virus and remained so even when longer virion-mAb interaction times were allowed. The V2q and V3 epitopes become available for binding by mAbs only when conformational changes were induced. This is expected for V3 epitopes, as V3 loops are suggested to be tucked underneath the V1V2 loops (105) and conformational changes such as movement or displacement of V1V2 can release the V3 loop allowing for its recognition by V3 targeting mAbs (106). Computational studies have also shown that the V3 loop can flicker in and out of its tucked position without disrupting the V1V2 organization at the apex (105, 107). However, these observations came from the simulation studies using the X-ray structure of the JR-FL SOSIP.559 trimer (PDB accession no. 5FYK), not virion associated Env, and therefore may or may not be applicable to all Env trimers. Even slight opening of the trimer apex can expose the highly immunogenic V3 loop (24). In such a case, we would expect to observe some level of binding to V3 mAbs. However, the V3 mAbs (2219, 2557) tested here did not show any binding even when the REJO virus and mAb interaction time were extended up to 75 min. The V3 epitopes also remained occluded in SF162 which is highly sensitive to neutralization by these V3 mAbs (71). The SF162 Env has historically been presumed to be in an open conformation (108–110), but based on the sensitivity of SF162 to neutralization by a family of mAbs that recognizes an SF162 type-specific quaternary epitope that bridges the V2 and V3 domains (111–114), it has been suggested that this Env actually exists predominantly in the closed conformation (115). This may explain why we observed masked V3 epitopes on SF162 in our assay. These data support the idea that V3 exposure varies among different HIV-1 isolates and was not applicable to the viruses tested here. Native-like Env trimers are the leading platform for HIV-1 vaccine design and several modifications have been made to reduce the Env metastability and exposure of the V3-loop (116–118). Understanding how the viruses retain the V3 in a tucked-in position may help these efforts.

The trimer apex epitopes on virions were inaccessible to the V2q mAbs that target these epitopes. Neutralization of HIV-1, or any other viral pathogen, minimally requires an initial encounter with an Ab. The PG9, PG16, and PGT145 mAbs can efficiently neutralize REJO virus (74). Thus, limited accessibility of these epitopes on virions was unexpected, particularly since V3 is shielded only when the native Env trimer is in the closed pre-fusion conformation: a conformation that is required for the binding of V2q mAbs (105, 119). It is known that vulnerable epitopes on HIV-1 particles are masked either conformationally or via the glycan shield (45, 120, 121). It is also known that due to this conformational masking, accessibility of epitopes to antibodies can be limited (122). The V2q mAbs are some of the most broad and potent bNAbs, making the V2q epitope region a major site of vulnerability (16). Thus, the finding that these functional and vulnerable bNAb Env epitopes are occluded on virions is not surprising. Considering that induction of bNAbs, including V2q type Abs, by immunization is a major goal of vaccine design, the data presented here helps us to better understand the strategies that can be employed to improve the presentation of these epitopes.

Our previously published study has shown that different mechanisms are involved in occluding the V2i and V3 epitopes (47). While V3 epitopes become accessible after engagement of CD4 receptor, this is not the case for V2i epitopes. Again, V2i epitopes such as those targeted by the 830A mAb are abundantly available on the virions, but both REJO and CMU06 viruses are resistant to neutralization by 830A when tested under standard 1 hour incubation assay conditions (47, 56, 74). Both viruses become neutralization sensitive if the virus-mAb incubation is allowed to continue to 24 hours prior to adding the target TZM.bl cells (56, 74). Thus, prolonged incubation time allows for conformational unmasking or allows Env breathing that may be transiently exposing the V2i and V3 epitopes. Allosteric changes induced by binding of V2i mAbs, but not prolonged interaction time with virus, allowed PG9 to bind to V2q epitopes on REJO virus. As both V2i and V2q mAbs bind V1V2 when V2C is in a β-strand conformation (46, 123), the mechanism by which the binding of V2i helped expose V2q remains unknown. Likewise, we observed no significant differences in neutralization of REJO virus by the V2q and CD4bs mAbs, when the neutralization assay omitted the commonly adopted one-hour pre-incubation step (85, 124–130). These data suggest that HIV-1 interaction with target cells causes changes that allow these bNAbs to bind to their target epitopes and neutralize virions. Glycan-glycan interactions are emerging as new class of high-affinity biomolecular interaction (131), and are shown to support the binding of SARS-CoV-2 to the host cell membrane (131–133). Similarly, it has been shown that glycan-glycan interaction between the HIV-1 Env N-linked glycans and the N-acetylglucosamine (GlcNAc) on the host cell glycocalyx can support virus adhesion to cells (96). Here we show that binding of PG9 to Env on REJO virions is detected in presence of 293T cells while negligible binding is observed when cells are not present, supporting that initial interaction with host cells may influence the Env organization exposing these epitopes. In addition, lectin-glycan interactions are also shown to impact virus transmission, virus attachment (134–136), and virus neutralization (99). Both ConA and GRFT lectins tested here can interact with Env to improve the exposure of the occluded epitopes. However, this effect is also dependent on the specificity of the lectin. Nonetheless, the evidence presented regarding how the occluded bNAb epitopes on virions may be unmasked, allowing the mAbs to bind to their epitopes and block infection, highlights an unexpected role for Env-cell interaction and a new perspective on the conformational dynamics of Env.

In summary, we show that only select Env epitopes are exposed on the HIV-1 surface, with V2i epitopes being the most abundant and potentially providing an explanation as to why Abs against these immunogenic epitopes are more readily elicited during infection (65, 66, 69, 70, 137–141). Pre-existing V2i antibodies can form immune complexes with HIV-1 Env and skew antibody elicitation to a more cross-reactive immune response (93, 94), suggesting a possible role of these Abs in influencing the quality of humoral immune response. We also show a previously unknown fact that interaction with host cells is required for bNAbs to be able to access their epitopes on virus particles. It will also be interesting to know if other viruses, for example SARS-CoV-2 and Influenza, have similar requirements concerning the interaction with host cells in order for the virus-specific neutralizing Abs to exert their effect. A greater understanding of epitope exposure on individual virions upon interaction with host cells would contribute greatly to understanding how humoral responses recognize different Env domains and how it may influence virus infection and antibody responses.

## Acknowledgments

The authors would like to thank Dr. Susan Zolla-Pazner and Ms. Xiaomei Liu for providing HIV-1-specific mAbs.

## Figure Legends

**Supplementary Figure 1.** Measurement of Env in virus preparations by Western blot. (A) Viruses produced in 293T cells, were concentrated (20X) by sucrose pelleting. Four μl of each 20X concentrated virus particle prep were lysed and analyzed by SDS-PAGE (4– 20%) and Western blot. An anti-gp120 MAb cocktail (V3: 391/95-D, 694/98-D, 2219, 2558; C2: 847-D, 1006-30D; C5: 450-D, 670-D) was used to quantitate the levels of Env associated with virions. REJO gp120 protein loaded at different concentrations was used as a standard. The band density of the REJO gp120 protein was used to generate the linear curve (B) and to calculate the amount of Env in each virus preparation.

**Supplementary Figure 2**. Histogram plots related to Fig 2 showing reactivity of serially diluted mAbs to REJO virus particles coupled to fluorescent microspheres. Virus coupled microspheres stained with biotin and PE alone were used to set the background staining (gray). Experiment was repeated to probe for binding of CD4-IgG, 17b and negative control mAb 3685 (Right panel)

**Supplementary Figure 3.** Histogram plots related to Fig 3 showing reactivity of serially diluted mAbs to SF162 virus particles coupled to fluorescent microspheres. Virus coupled microspheres stained with biotin and PE alone were used to set the background staining (gray).

**Supplementary Figure 4.** Representative histogram plots related to Fig 4 showing reactivity of mAbs to CMU06 virus particles coupled to fluorescent microspheres. Binding was detected with streptavidin-PE and fluorescent intensity (MFI) are shown. Virus coupled microspheres stained with biotin and PE alone were used to set the background staining (gray).

**Supplementary Figure 5.** Histogram plots related to Fig 5A showing reactivity of mAbs to REJO Env expressed on cell surface. Replicates from one representative experiment are shown. Binding was detected with streptavidin-PE. Transfected cells stained with biotin and PE alone were used to set the background MFI (gray). MAb 3685 was used as negative control.

**Supplementary Figure 6.** Histogram plots related to Fig 5B showing reactivity of mAbs to CMU06 Env expressed on cell surface. Triplicates are shown. Binding was detected with streptavidin-PE. Transfected cells stained with biotin and PE alone were used to set the background MFI (gray). MAb 3685 was used as negative control.

**Supplementary Figure 7.** Histogram plots related to Fig 5C showing reactivity of mAbs to SF162 Env expressed on cell surface. Triplicates for all antibodies and duplicates for CD4-IgG and PGT151 are shown. Binding was detected with streptavidin-PE. Transfected cells stained with biotin and PE alone were used to set the background MFI (gray). MAb 3685 was used as negative control.

**Supplementary Figure 8.** Representative histogram plots related to Fig 6 showing time-dependent changes in reactivity of mAbs. Microspheres coupled to REJO virus particles were treated with mAbs at 37°C for various time from 0 to 75 min. Binding was detected with streptavidin-PE.

**Supplementary Figure 9.** Representative histogram plots related to Fig 7 showing changes in reactivity of PG9 and 2219 mAbs by different antibodies. Microspheres coupled with REJO virus were incubated with five-fold titrated antibodies followed by biotinylated V2q mAb PG9 (25 μg/ml). Binding was detected with streptavidin-PE.

**Supplementary Figure 10. Gating strategy** (A) **and histograms related to Fig 10**. Plots showing binding of mAbs to virions in presence of (B) 293T cells transfected to express hCD4 and (C) un-transfected 293T cells. (D) Histograms showing cell surface expression of human CD4 293T cells. Cells expressing hCD4 are shown in blue and un-transfected cells are shown in gray.

**Supplementary Figure 11.** Representative histogram plots related to Fig 11 showing changes in reactivity of mAbs by Concanavalin A (ConA).

## Institutional Review Board Statement

“Not applicable”.

## Informed Consent Statement

“Not applicable”.

## Data Availability Statement

All available data are presented in the article.

