## Supplementary Figures for "Broadly Neutralizing Antibody Epitopes on HIV-1 Particles are exposed after Virus Interaction with Host Cells"

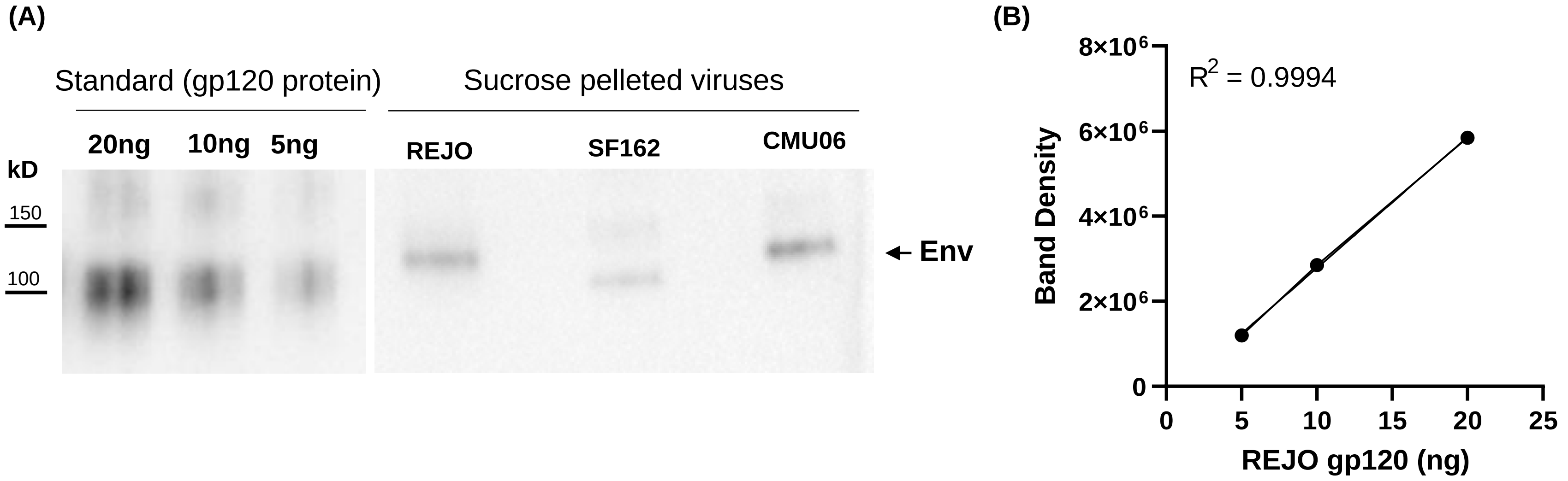


**Supplementary Figure 1. Measurement of Env in virus preparations by Western blot.** (A) Viruses produced in 293T cells, were concentrated (20X) by sucrose pelleting. Four µl of each 20X concentrated virus particle prep were lysed and analyzed by SDS-PAGE (4–20%) and Western blot. An anti-gp120 MAb cocktail (V3: 391/95-D, 694/98-D, 2219, 2558; C2: 847-D, 1006-30D; C5: 450-D, 670-D) was used to quantitate the levels of Env associated with virions. REJO gp120 protein loaded at different concentrations was used as a standard. The band density of the REJO gp120 protein was used to generate the linear curve (B) and to calculate the amount of Env in each virus preparation.

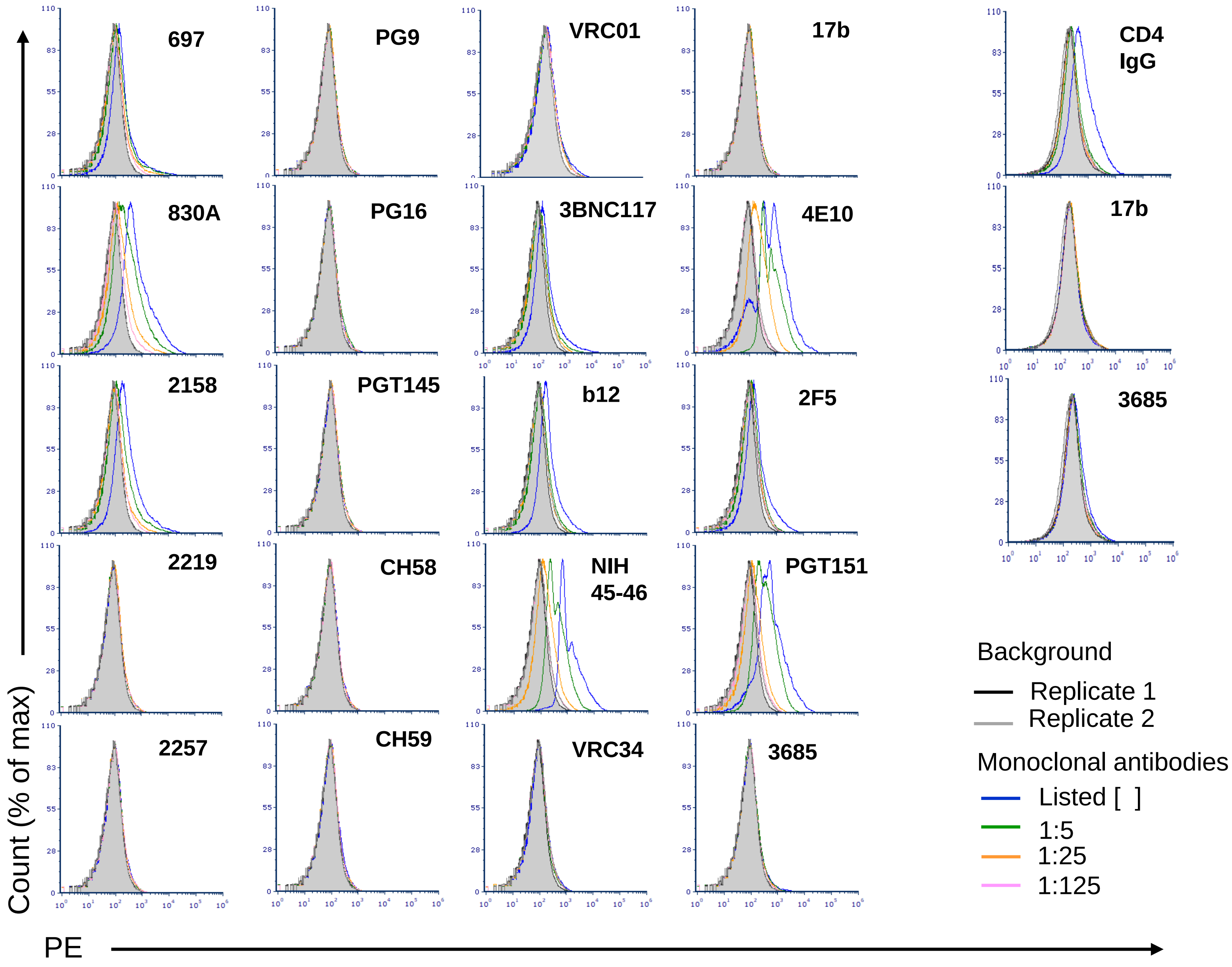


**Supplementary Figure 2. Histogram plots related to Fig 2 showing reactivity of serially diluted mAbs to REJO virus particles coupled to fluorescent microspheres.** Virus coupled microspheres stained with biotin and PE alone were used to set the background staining (gray). Experiment was repeated to probe for binding of CD4-IgG, 17b and negative control mAb 3685 (Right panel)

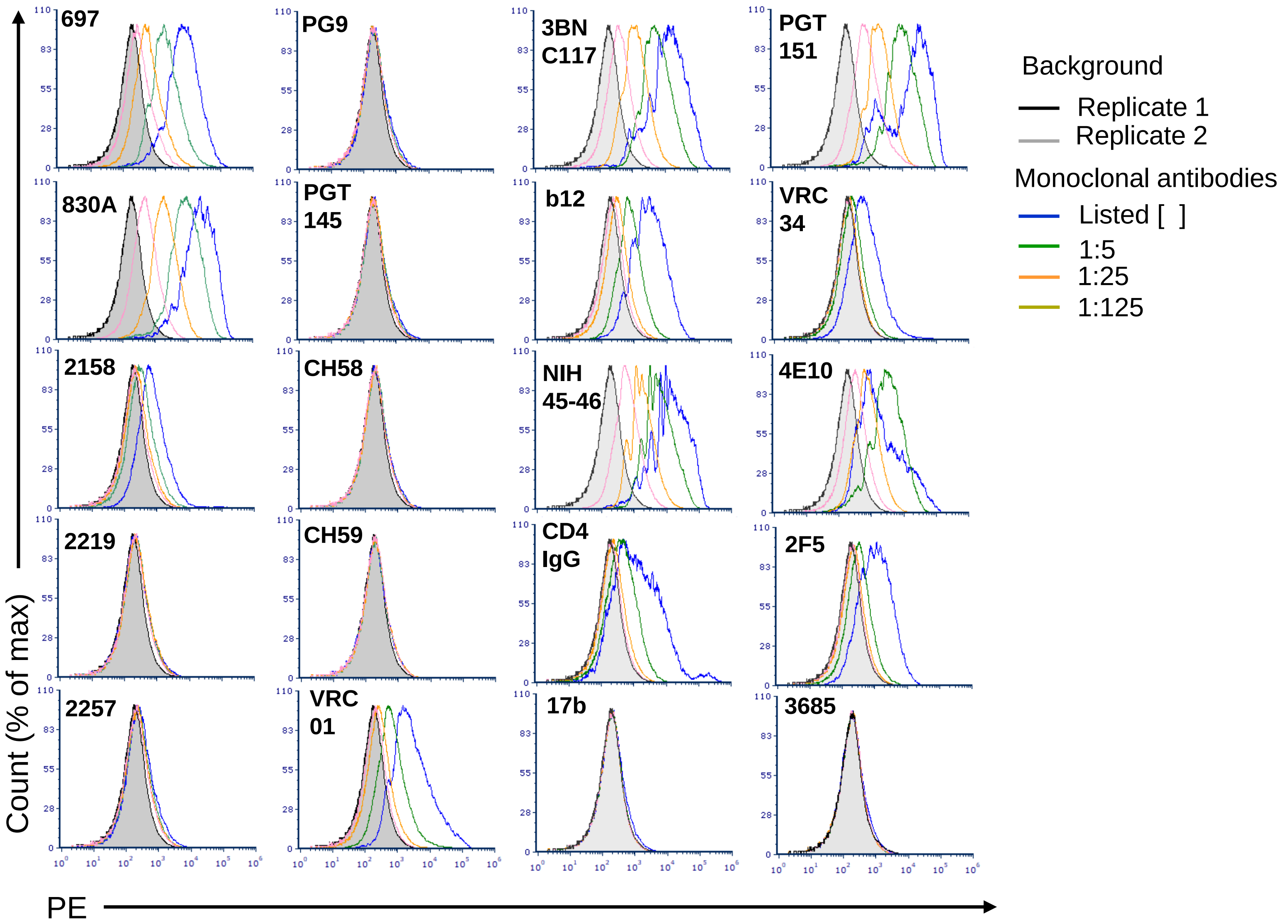


**Supplementary Figure 3. Histogram plots related to Fig 3 showing reactivity of serially diluted mAbs to SF162 virus particles coupled to fluorescent microspheres.** Virus coupled microspheres stained with biotin and PE alone were used to set the background staining (gray).

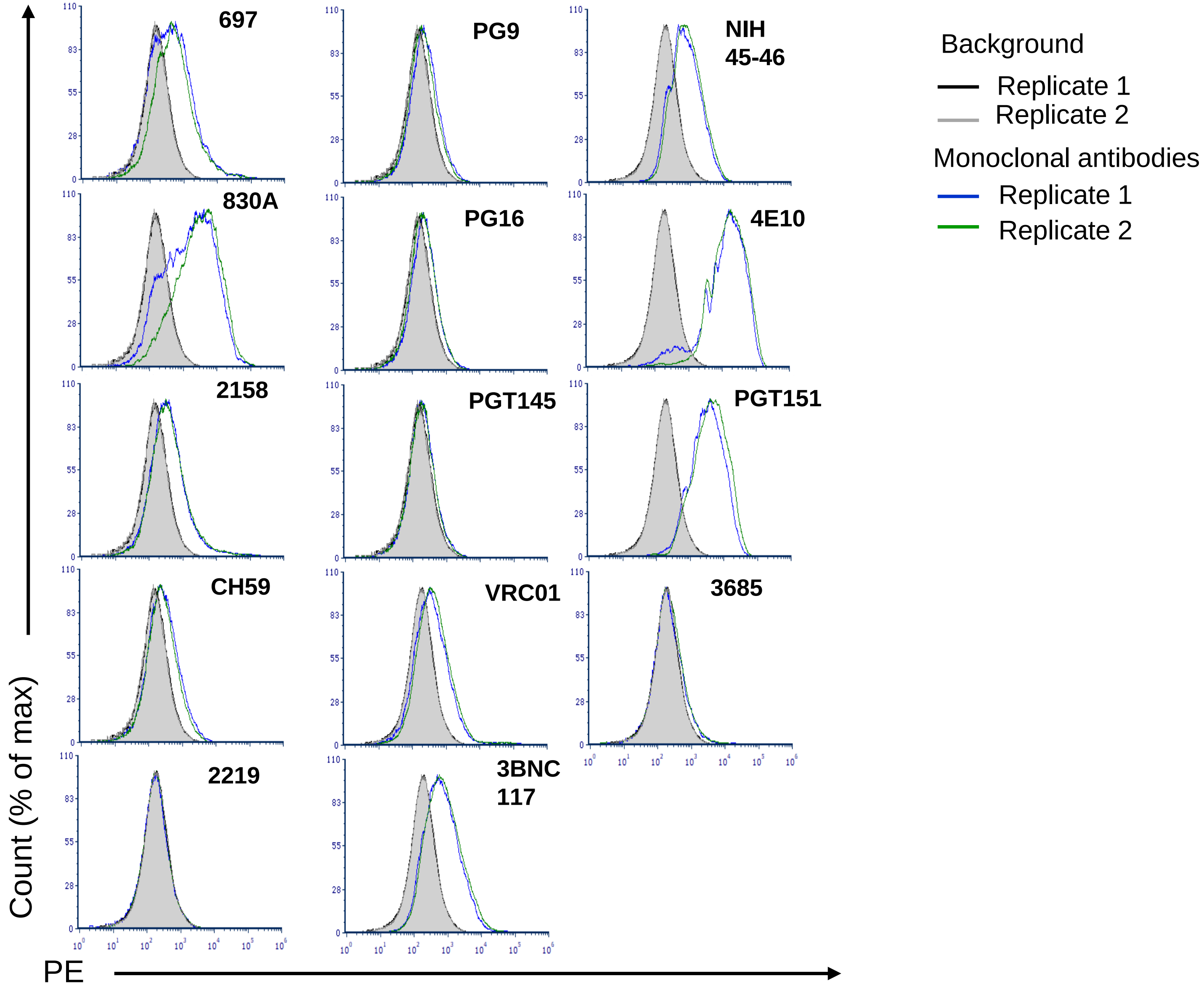


**Supplementary Figure 4. Representative histogram plots related to Fig 4 showing reactivity of mAbs to CMU06 virus particles coupled to fluorescent microspheres.** Binding was detected with streptavidin-PE and fluorescent intensity (MFI) are shown. Virus coupled microspheres stained with biotin and PE alone were used to set the background staining (gray).

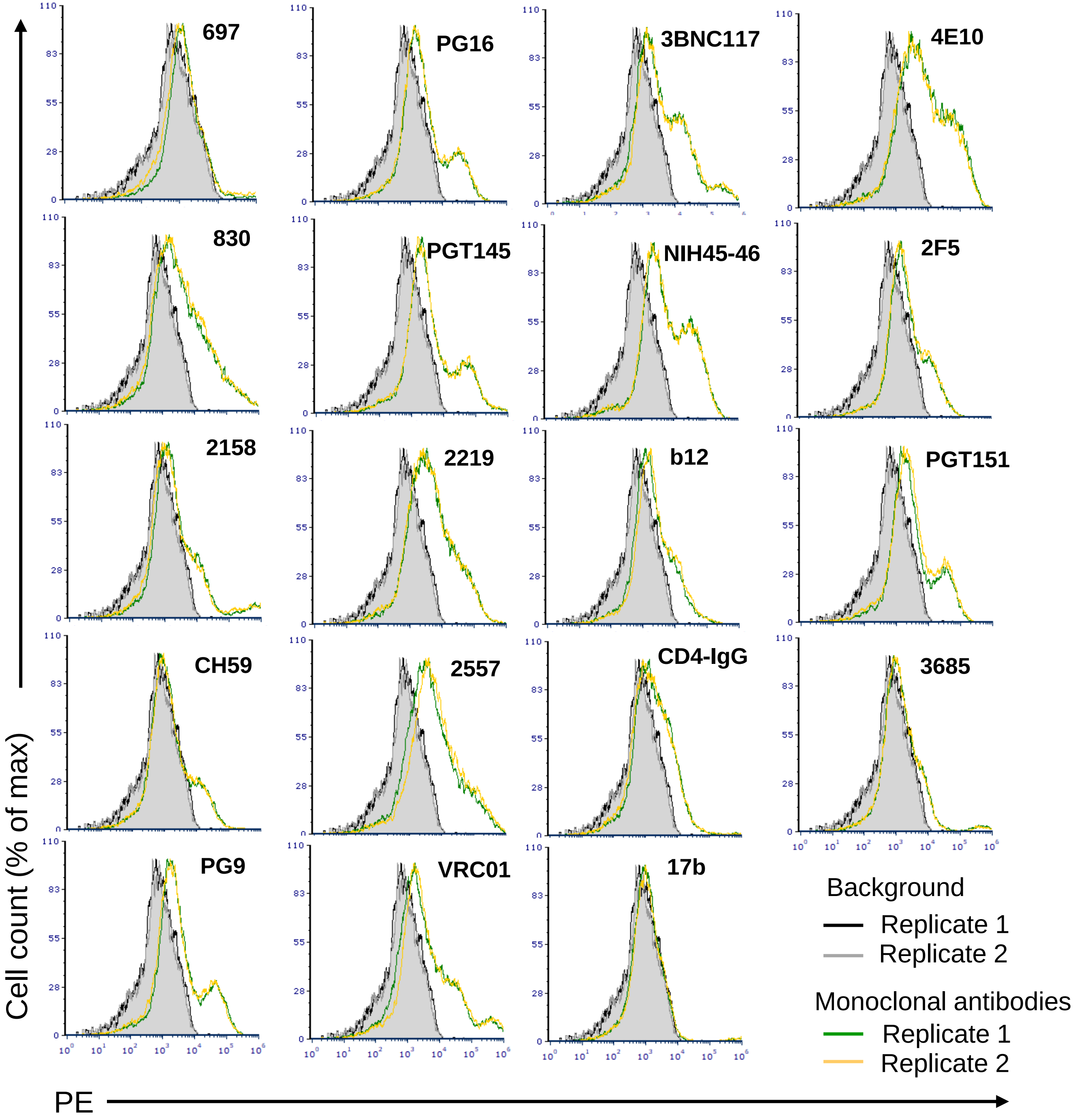


**Supplementary Figure 5. Histogram plots related to Fig 5A showing reactivity of mAbs to REJO Env expressed on cell surface. Replicates from one representative experiment are shown.** Binding was detected with streptavidin-PE. Transfected cells stained with biotin and PE alone were used to set the background MFI (gray). MAb 3685 was used as negative control.

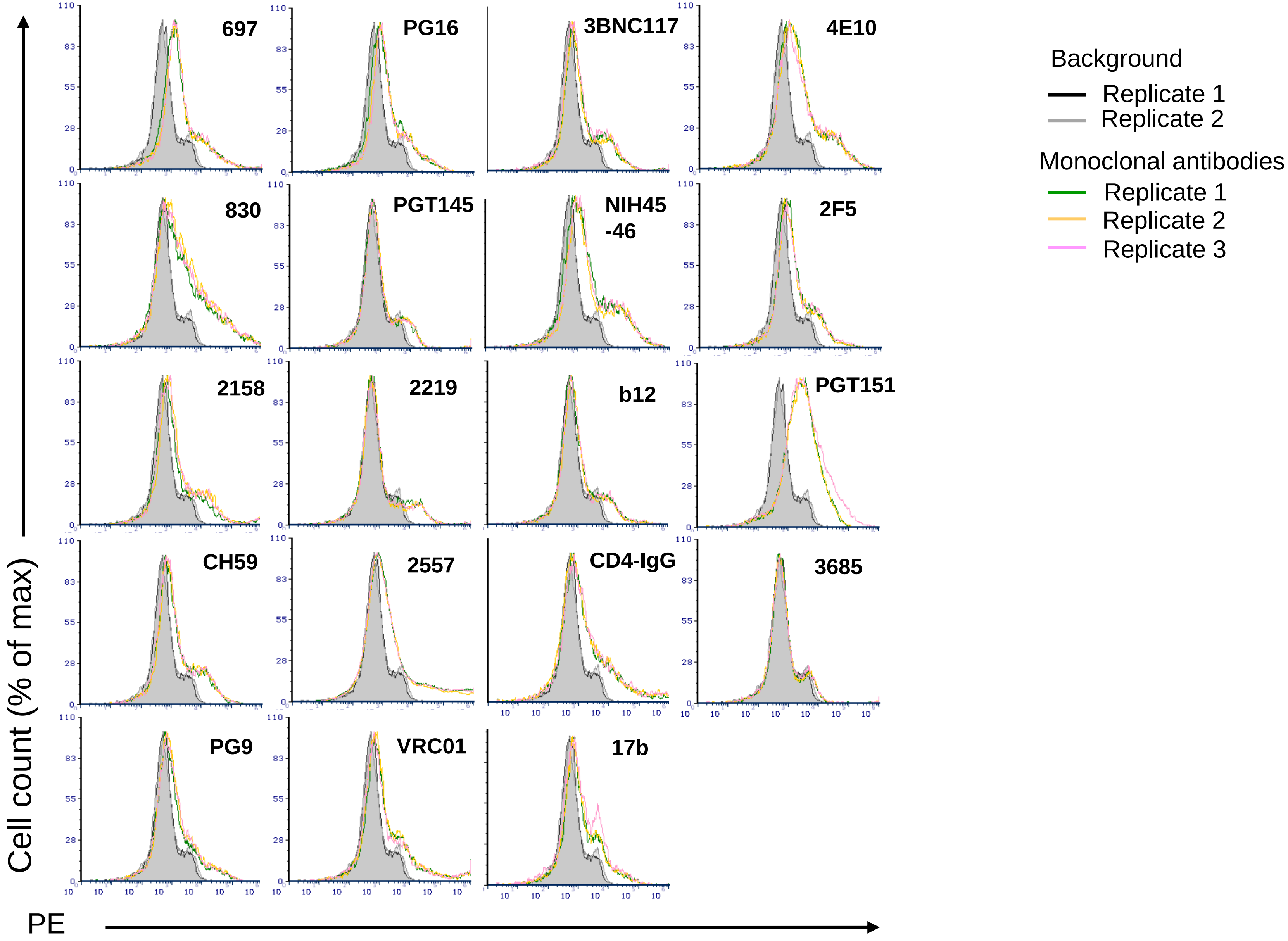


**Supplementary Figure 6. Histogram plots related to Fig 5B showing reactivity of mAbs to CMU06 Env expressed on cell surface.** Triplicates are shown. Binding was detected with streptavidin-PE. Transfected cells stained with biotin and PE alone were used to set the background MFI (gray). MAb 3685 was used as negative control.

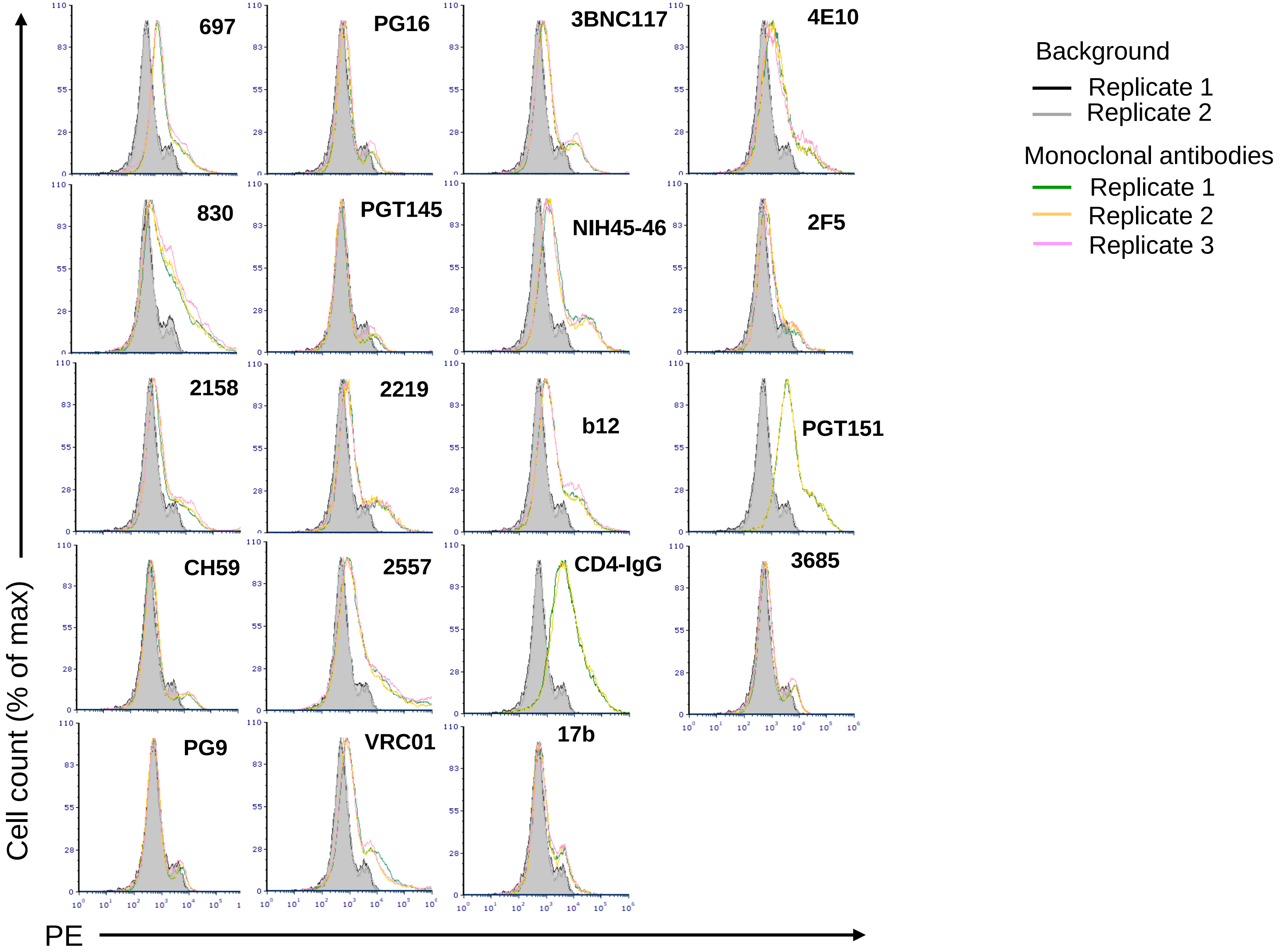


**Supplementary Figure 7. Histogram plots related to Fig 5C showing reactivity of mAbs to SF162 Env expressed on cell surface. Triplicates for all antibodies and duplicates for CD4-IgG and PGT151 are shown.** Binding was detected with streptavidin-PE. Transfected cells stained with biotin and PE alone were used to set the background MFI (gray). MAb 3685 was used as negative control.

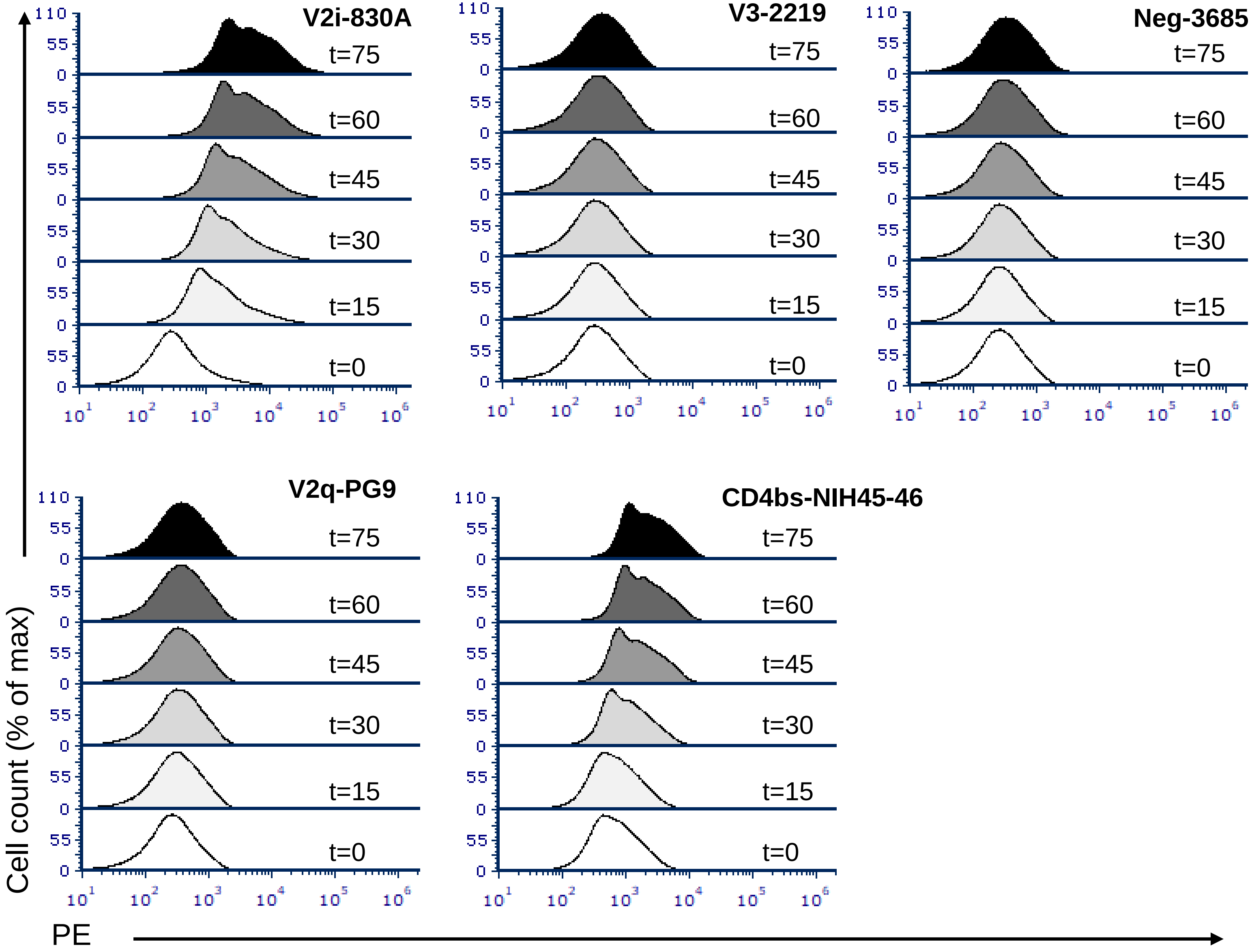


**Supplementary Figure 8. Representative histogram plots related to Fig 6 showing time-dependent changes in reactivity of mAbs.** Microspheres coupled to REJO virus particles were treated with mAbs at 37°C for various time from 0 to 75 min. Binding was detected with streptavidin-PE.

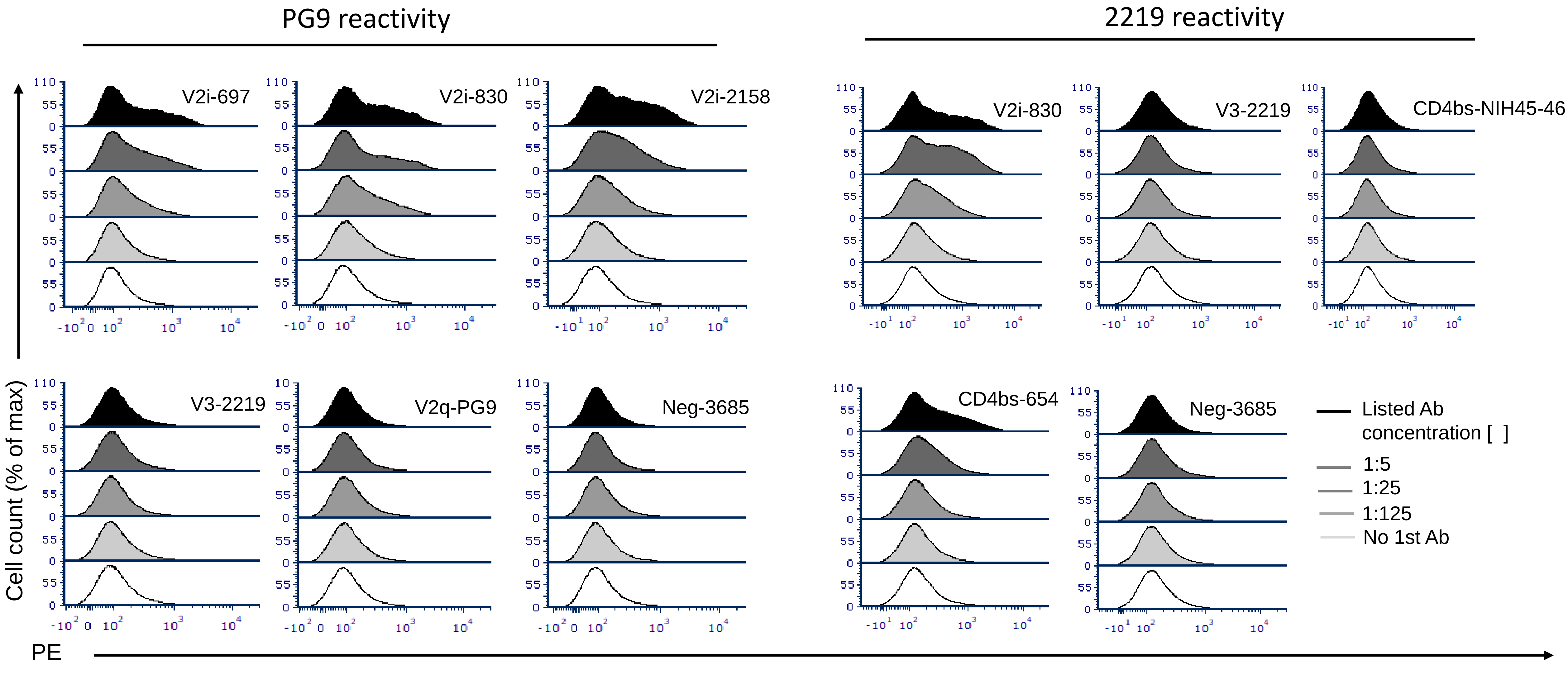


**Supplementary Figure 9. Representative histogram plots related to Fig 7 showing changes in reactivity of PG9 and 2219 mAbs by different antibodies.** Microspheres coupled with REJO virus were incubated with five-fold titrated antibodies followed by biotinylated V2q mAb PG9 (25 µg/ml). Binding was detected with streptavidin-PE.

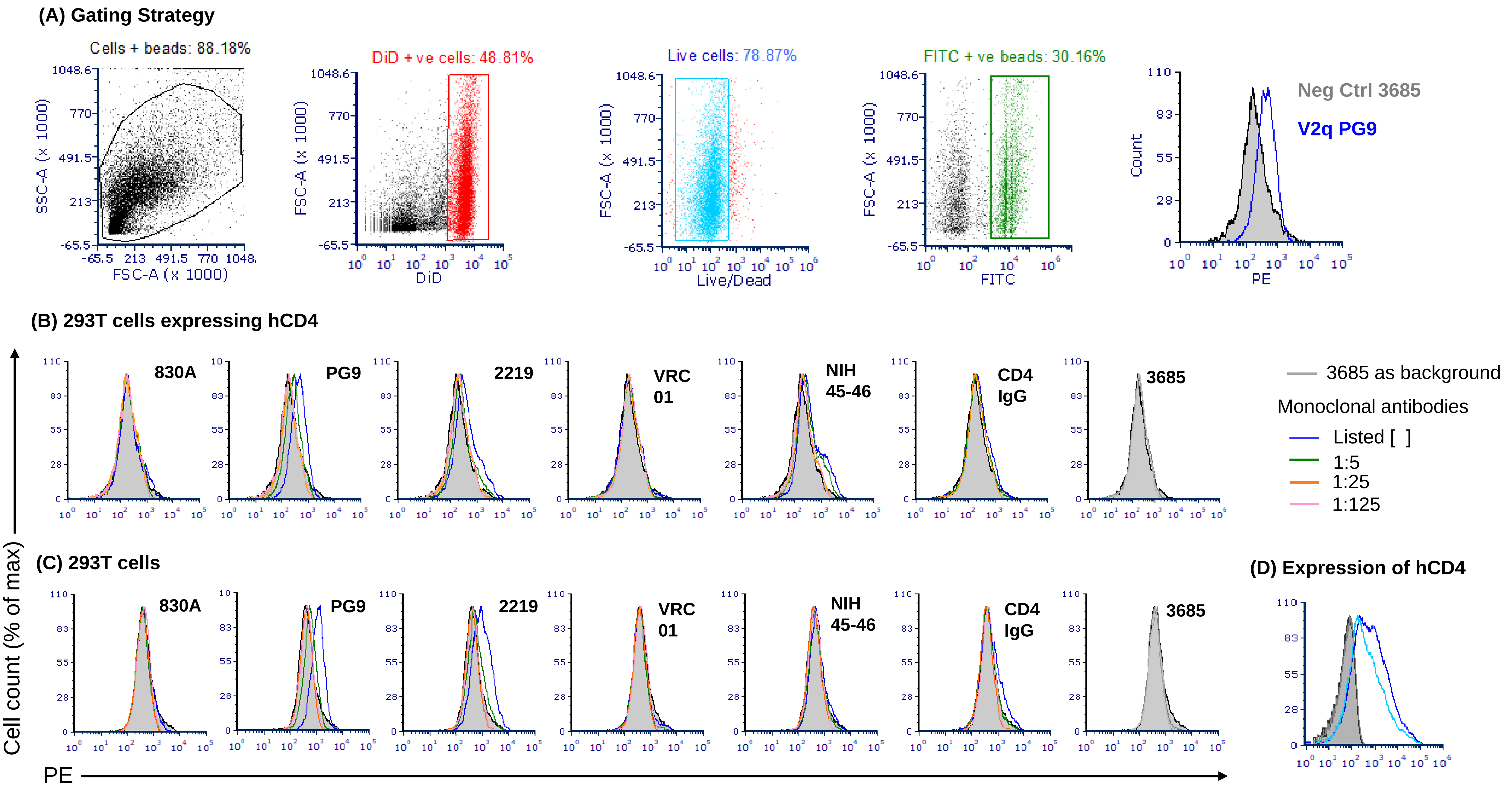


**Supplementary Figure 10. Gating strategy** (A) **and histograms related to Fig 10.** Plots showing binding of mAbs to virions in presence of (B) 293T cells transfected to express hCD4 and (C) un-transfected 293T cells. (D) Histograms showing cell surface expression of human CD4 293T cells. Cells expressing hCD4 are shown in blue and un-transfected cells are shown in gray.

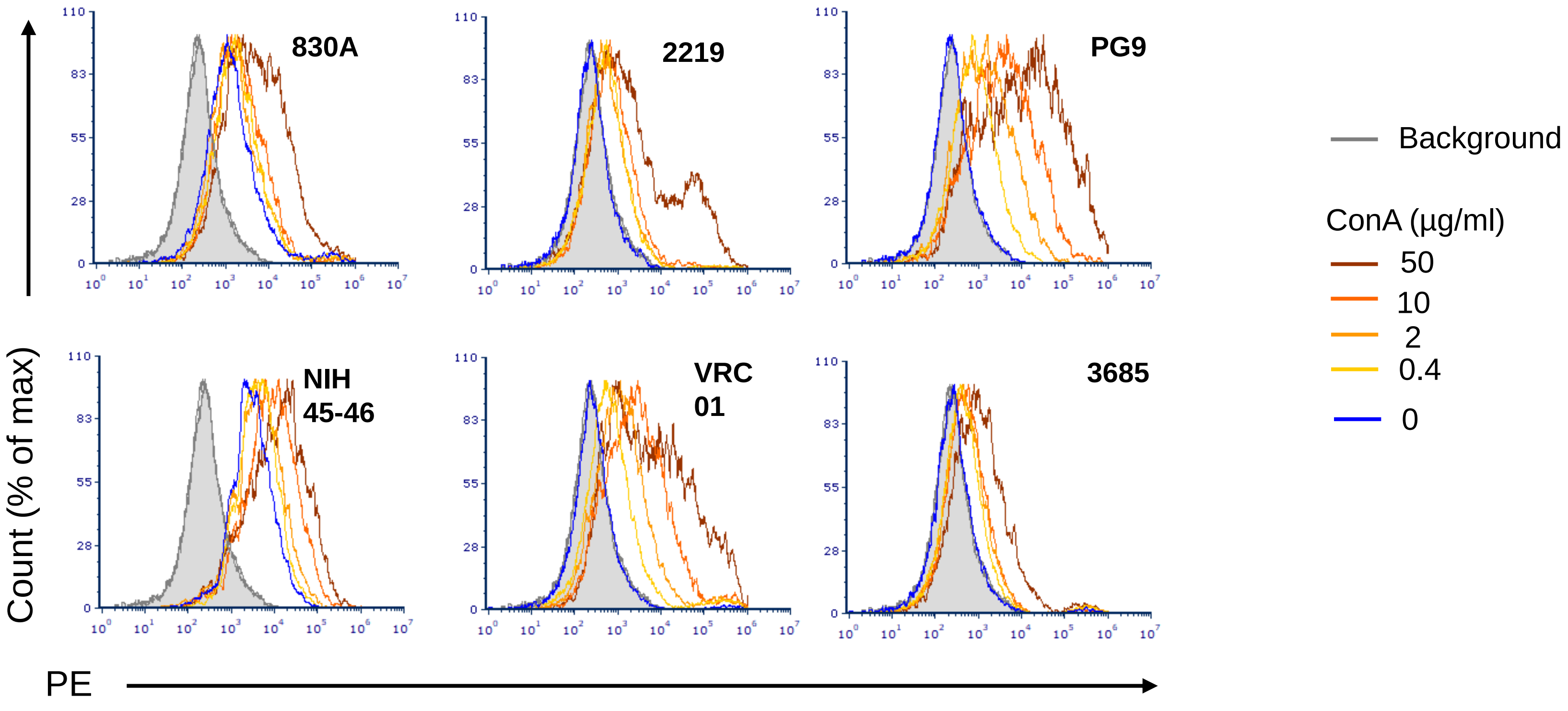


**Supplementary Figure 11.** Representative histogram plots related to Fig 11 showing changes in reactivity of mAbs by Concanavalin A (ConA).
